# Repeat-based holocentromeres of the woodrush *Luzula sylvatica* reveal new insights into the evolutionary transition from mono- to holocentricity

**DOI:** 10.1101/2024.04.05.588299

**Authors:** Yennifer Mata-Sucre, Marie Krátká, Ludmila Oliveira, Pavel Neumann, Jiří Macas, Veit Schubert, Bruno Huettel, Eduard Kejnovský, Andreas Houben, Andrea Pedrosa-Harand, Gustavo Souza, André Marques

## Abstract

Although the centromere is restricted to a single region of the chromosome in most studied eukaryotes, members of the rush family (Juncaceae) harbor either monocentric (*Juncus)* or holocentric (*Luzula*) chromosomes. This provides an opportunity to study the evolutionary mechanisms involved in the transition to holocentricity. Here by combining chromosome-scale genome assembly, epigenetic analyses, immuno-FISH, and super-resolution microscopy, we report the occurrence of repeat-based holocentromeres in *L. sylvatica*. We found an irregular distribution of genes, centromeric units, and most repeats along the chromosomes. We determined the centromere function predominantly associated with two satellite DNA repeats, *Lusy1* and *Lusy2* of 124– and 174-bp monomer length, respectively, while CENH3 also binds satellite-free gene-poor regions. Comparative repeat analysis revealed that *Lusy1* is present in most *Luzula* species, suggesting a conserved centromere role of this repeat. Synteny between *L. sylvatica* (*n* = 6) and *J. effusus* (*n* = 21) genomes further evidenced a chromosome number reduction in *Luzula* derived from multiple chromosome fusions of ancestral *J. effusus*-like chromosomes. We propose that the transition to holocentricity in *Luzula* involves: (i) fusion of small chromosomes resembling *Juncus*-like centromeres; (ii) expansion of atypical centromeric units; and (iii) colonization of satellite DNA for centromere stabilization.

## Introduction

Centromeres are specialized chromosomal regions that recruit kinetochore proteins and mediate spindle microtubule binding to ensure correct chromosome segregation during mitosis and meiosis (Talbert and Henikoff 2020; Schubert *et al*. 2020). Most taxonomic groups have chromosomes with a size-restricted centromeric domain confined to the primary chromosome constriction, i.e., they are monocentric (Wong *et al*. 2020). However, holocentric chromosomes lack a primary constriction and exhibit molecular and epigenetic features that allow kinetochore proteins and microtubules to bind extensively along the chromosomes (Heckmann *et al*. 2013; Schubert *et al*. 2020). Therefore, holocentric chromosomes can tolerate large-scale rearrangements, such as chromosome fusions and fissions, because the rearranged chromosomes can maintain kinetochore activity, avoiding segregation problems and preserving essential genetic information during cell divisions (Burchardt *et al*. 2020; Hofstatter *et al*. 2022; Escudero *et al*. 2023; Mata-Sucre *et al*. 2024).

Remarkably, holocentric chromosomes have evolved repeatedly in animals and plants (Melters *et al*. 2012; Escudero *et al*. 2016; Senaratne *et al*. 2022). The lack of conclusive evidence pointing to reversions to monocentricity in any eukaryotic lineage and the sporadic distribution of holocentric versus monocentric organisms support the unidirectional transition to holocentricity (Escudero *et al*. 2016). Numerous evolutionary models have been proposed to explain the emergence of holocentricity from monocentric ancestors, which include alterations/loss/emergence of kinetochore genes or centromeric repetitive sequences during the process (Drinnenberg *et al*. 2014; Senaratne *et al*. 2021; Neumann *et al*. 2023). The causes of the transitions, however, remain unclear, mainly because only a few holocentric species have been studied and because most holocentric groups evolved a long time ago, making the factors involved in the transition difficult to determine (Neumann *et al*. 2023).

In most plants, functional centromeres are epigenetically specified by the centromeric histone H3 variant (CENH3). CENH3 binding regions (hereafter CENH3 domains) in monocentric chromosomes are typically associated with extended arrays of tandemly repeated sequences (satellite DNA), which are usually highly divergent and fast evolving (Allshire and Karpen 2008; Plohl *et al*. 2014; Ávila Robledillo *et al*. 2020; Šatović-Vukšić and Plohl 2023). Although the role of these repeats in centromere function has not yet been fully elucidated, several possible advantages of centromeric repeats have been proposed. Satellites might have favorable monomer lengths stabilizing CENH3 nucleosome positioning or contain specific sequences, such as short dyad symmetries, forming non-B-DNA structures possibly aiding CENH3 nucleosome loading through interaction with CENH3 chaperone HJURP (Hobza *et al*. 2006; Kasinathan and Henikoff 2018; Naughton and Gilbert 2020; Talbert and Henikoff 2020). Nevertheless, only a few holocentric species with centromeric repeats have been characterized so far. *Rhynchospora* Vahl. (Cyperaceae) holocentromeres are mainly composed of a 172-bp satellite called *Tyba*, evenly distributed along the chromosomes in ∼20kb domains and specifically colocalizing with CENH3 (Marques *et al*. 2015; Hofstatter *et al*. 2022; Castellani *et al*. 2024). In *Chionographis japonica* (Willd.) Maxim. (Melanthiaceae), few large (∼2Mb) CENH3-positive domains are associated with satellite arrays of 23– and 28-bp-long monomers (Kuo *et al*. 2023). Similarly, in mulberry (*Morus notabilis*) few CENH3-positive domains are associated with satellite arrays of 82-bp-long monomers (Ma *et al*. 2023). Furthermore, the *Meloidogyne incognita* root-knot nematode is so far the only holocentric animal to show a 19-bp sequence box conserved within diverse centromeric satellites associated with holocentromere function (Despot-Slade *et al*. 2021).

Juncaceae Juss. (rushes/woodrushes), the sister family of Cyperaceae (sedges), is a cosmopolitan family comprising ∼473 species (POWO 2024). *Juncus* L. (rush) and *Luzula* DC. (woodrush) represent the largest genera in the family with 332 and 124 species, respectively ( POWO 2024). An interesting feature of this family is its variation in centromeric organization and chromosomal structure, making it an ideal model to address hypotheses about evolutionary processes during centromere development. Although historically the entire Juncaceae family was thought to be holocentric, cytogenetic and genomic studies revealed that six different *Juncus* species are monocentric (Guerra *et al*. 2019; Hofstatter *et al*. 2022; Mata-Sucre *et al*. 2023; Dias *et al*. 2024). Recent chromatin immunoprecipitation sequencing revealed that *J. effusus* has repeat-based and CENH3-associated monocentromeres, consisting mainly of two tandem repeat families underlying one or up to three spaced cores of CENH3-enriched regions per chromosome (Dias *et al*. 2024). On the other hand, *Luzula* species have been up till now characterized as holocentric without specific centromeric repeats (Kuta *et al*. 2004; Nagaki *et al*. 2005; Heckmann *et al*. 2011, 2013; Bozek *et al*. 2012). Although in *Luzula* species, a 178 bp satellite sharing sequence similarity with the rice centromeric repeat was discovered (Haizel *et al*. 2005), it is uncertain whether this satellite plays a centromeric role and the lack of a reference genome has made detailed studies of *Luzula* centromeres challenging.

Here we performed a comprehensive (epi)genomic characterization of the chromosome-scale genome of the woodrush *L. sylvatica*, focusing on its holocentromere organization, repetitive fraction and genome evolution. We show that *L. sylvatica* has a unique repeat-based centromere organization distinct from previously described holocentric species. Comparative genomic repeat profiles of 13 *Luzula* species revealed likely conservation of repeat-based holocentromeres in the genus, except for *L. elegans*. Further comparative genomics analysis between *J. effusus* and *L. sylvatica* revealed footprints of extensive monocentric chromosome fusions that potentially played an important role in the transition to holocentricity in the lineage.

## Results

### Chromosome-scale genome assembly reveals the repeat-based holocentromeres of *L. sylvatica*

We estimated a genome size of 1C = 476 Mb for *L. sylvatica* (2*n* = 12) based on k-mer frequencies (**Fig. S1**), and assembled a chromosome-scale reference genome sequence integrating PacBio HiFi reads and a Hi-C chromatin interaction dataset available in www.darwintreeoflife.org (**Fig. 1, Fig. S1**). The *de novo* genome assembly of *L. sylvatica* generated 1,010 contigs totaling 516.08 Mb with a GC content of 33.01 %, N50 of 7.6 Mb, and BUSCO completeness of 93.01% (**Fig. 1a-b, Table S1**). Six pseudomolecules were obtained by Hi-C scaffolding, with a total of 468.44 Mb and an N50 of 78.95 Mb (**Fig. 1c, Table S1**). Similar to holocentric beak-sedges (Hofstatter *et al*. 2022), as the concept of chromosome arms does not apply to holocentric species, we observed no large-scale compartmentalization or telomere-to-centromere axis, as evidenced by the chromatin configuration capture (Hi-C) contact matrix (**Fig. 1c**). Immunolabelling of CENH3 on *L. sylvatica* mitotic cells confirmed the holocentricity of its chromosomes (**Fig. 1d**). The assembly was annotated concerning major genomic sequence types, including genes, tandem repeats and transposable elements (**Fig 1e, Fig. S2**). A dispersed but structurally heterogeneous distribution of sequences along all pseudomolecules was observed, with interstitial regions highly enriched by tandem repeats and lacking genes and transposable elements (**Fig 1e, Fig. S2**).

**Fig. 1:**
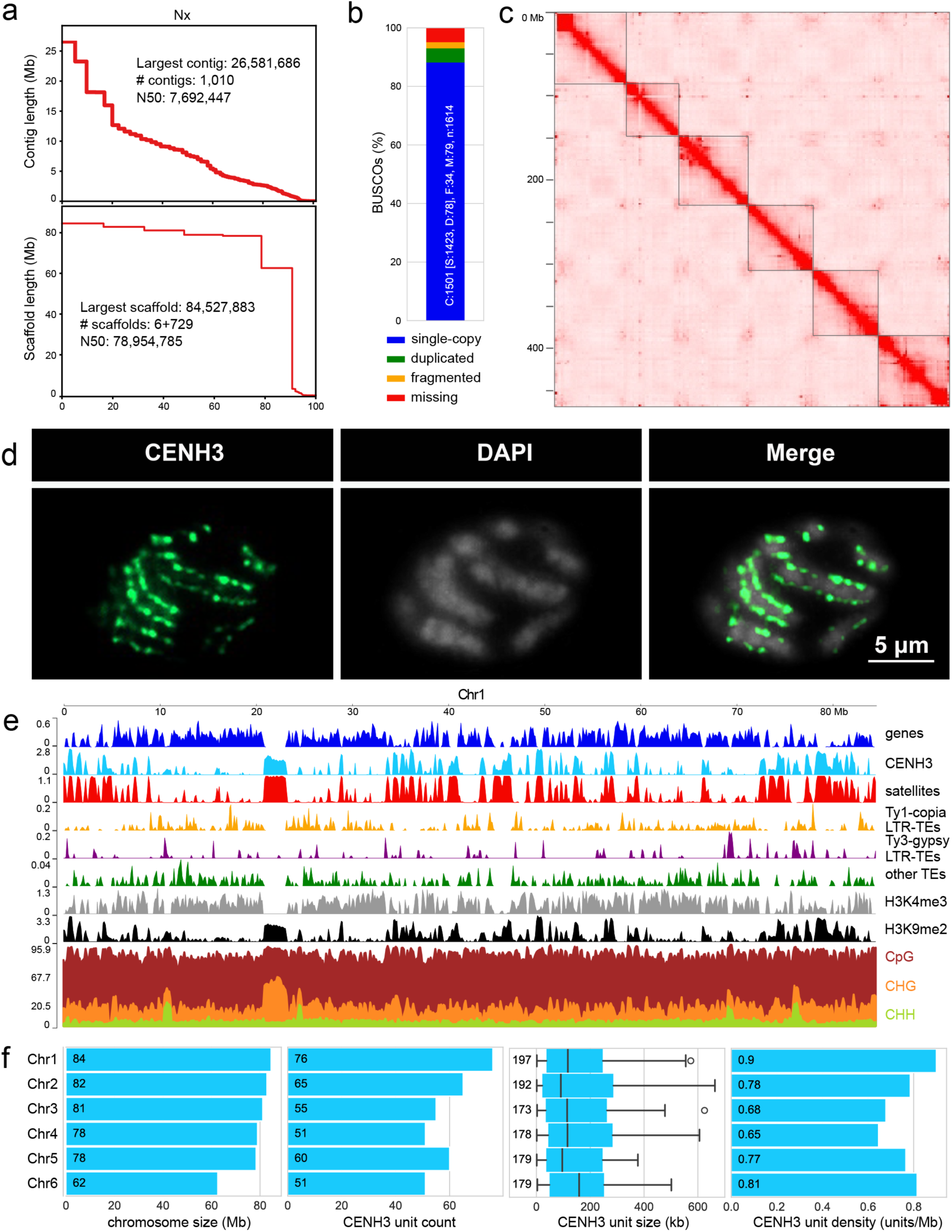
Genome assembly and annotation of *L. sylvatica*. **(a)** Statistics of the *L. sylvatica* genome assembly (top) and the final scaffolding (bottom). **(b)** BUSCO assessment for completeness of genic space with the viridiplantae_odb10 dataset, using the entire genome assembly. **(c)** Intra and inter-chromosome contact matrices of *L. sylvatica*. Color intensity represents contact frequency. Dark lines mark chromosomal boundaries. Boxes along the diagonal represent interactions within the same chromosome (cis), as expected for holocentric chromosomes. **(d)** Immunostaining of *L. sylvatica* centromeres using CENH3 antibodies. **(e)** *L. sylvatica* Chr1 detailed view showing dispersed distribution of main genomic features: CENH3, gene, tandem repeat, dispersed repeat and eu-heterochromatin marks densities, and methylation; as typical for holocentric chromosomes. Bin sizes of 100kb. The distribution of features on all chromosomes is reported in Fig. S2. **(f)** Size of chromosomes, the number of CENH3 domains, their size (<700 kb shown), and density (units/Mb).

To identify and characterize the centromeres, as well as eu– and heterochromatin regions of the *L. sylvatica* genome, we further performed chromatin immunoprecipitation with sequencing (ChIP-seq) for CENH3, H3K4me3, and H3K9me2, along with DNA methylation sequencing (**see Online Methods, Fig. 1e-f**). We detected 358 CENH3 domains distributed across the entire length of all chromosomes (**Fig. 1e-f, S2**). Considering that one CENH3 domain is equivalent to one centromeric unit, we observed an average of 0.76 units/Mb (range 0.64-0.90 units/Mb) or 60 units (range 51-76 units) per chromosome with an average unit length of 183 kb (range 174-197 kb; **Fig. 1f**). Additionally, histone modification marks H3K4me3 and H3K9me2 were intermixed along the chromosomes without a chromosome-wide pattern (**Fig. 1e, Fig. S2**).

The annotation of the genome repetitive fraction, which represents ∼59% of the genome, was based on DANTE and TAREAN (**Table 1, see Methods**). Most of these corresponded to satellite DNA sequences with six families representing 35.31 % of the genome, where the CL1 and CL2 clusters correspond to the most abundant satellite DNAs with 25.10 % and 7.06 %, respectively (**Table S2 and Fig. S3**). CL1 is a 124-bp satellite, named hereafter as *Lusy1* (**Table S2 and Fig. S3b**). CL2 is a satellite consisting of two variants of 174 and 175 bp sharing 62% similarity (hereafter referred to as *Lusy2*; **Table S2 and Fig. S3**), and only 30% similarity to *Lusy1*. The other five satellite DNAs have monomers with 31 to 182 bp, amounting to less than ∼2 % in the genome each (**Fig. 2a and Table S2**). Retrotransposon elements were less abundant than satellites, making up 18% of the genome (**Table 1)**. LTR retrotransposons of the *Ty1-copia* superfamily were the most represented with the *Angela* lineage being the most abundant (10.15 %; **Table 1**, **Fig. 1d**).

**Fig. 2:**
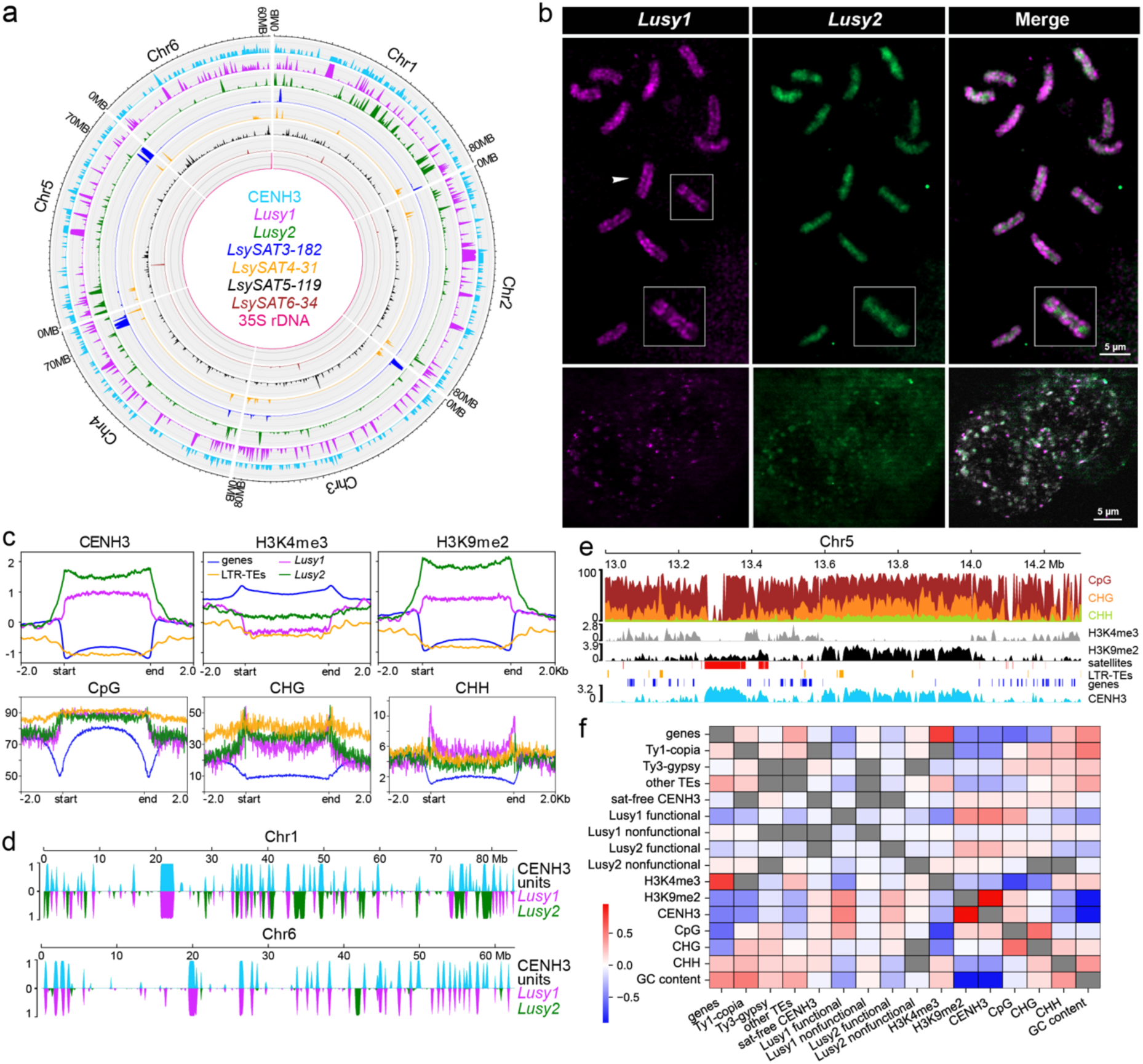
*L. sylvatica* holocentromeres are based on satellite repeats. **(a)** Circos distribution of the main classes of tandem sequence types and CENH3 domains with a 300kb sliding window. **(b)** FISH showing discontinuous linear-like spreading of the *Lusy1* (magenta) and disperse pattern of *Lusy2* (green) repeats in the nucleus and metaphase chromosomes. **(c)** Metaplots showing the enrichment of CENH3, H3K4me3, H3K9me2, CpG, CHH, and CGH from the start and end of different types of sequences: genes (blue), LTR transposable elements (yellow), *Lusy1* (magenta) and *Lusy2* repeats (green). ChIP-seq signals are shown as log2 (normalized RPKM ChIP/input). Methylation signals are shown as a percentage of methylated bases in each (CpG, CHG, CHH) context. **(d)** Proportion of CENH3 domains (light blue), *Lusy1* (magenta) and *Lusy2* (green) arrays in 100kb windows. Distribution on all chromosomes is reported in **Fig. S4**. **(e)** Close-up view of a genomic locus showing both *Lusy* satellite-based and satellite-free CENH3 domains. Distribution on all chromosomes is reported in **Fig. S5**. **(f)** Correlogram of genomic features in 100kb windows (n = 4694). Gray fields indicate values on the diagonal and non-significant values of the Spearman coefficient after multiple-testing correction (see Methods).

**Table 1.**
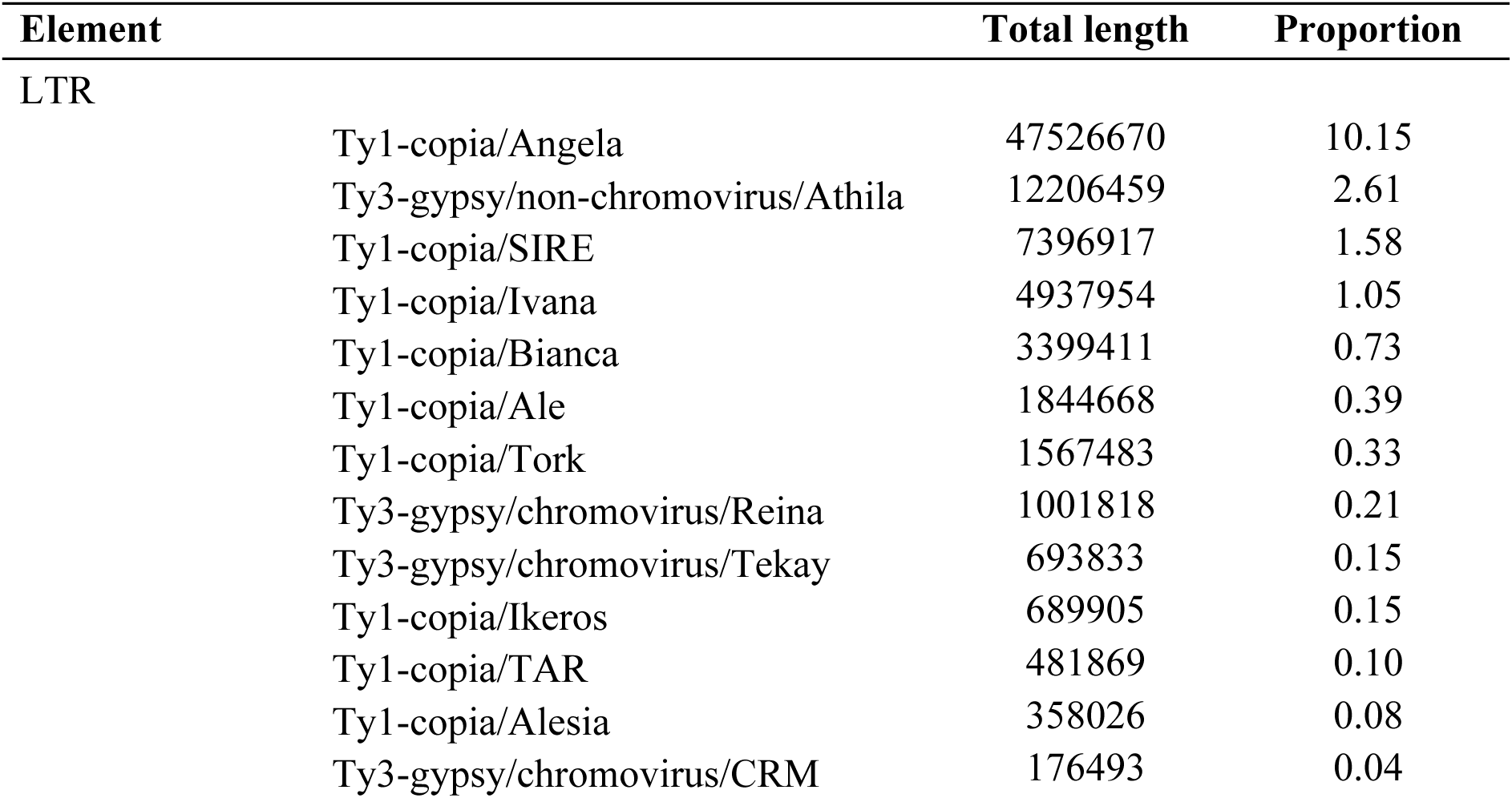
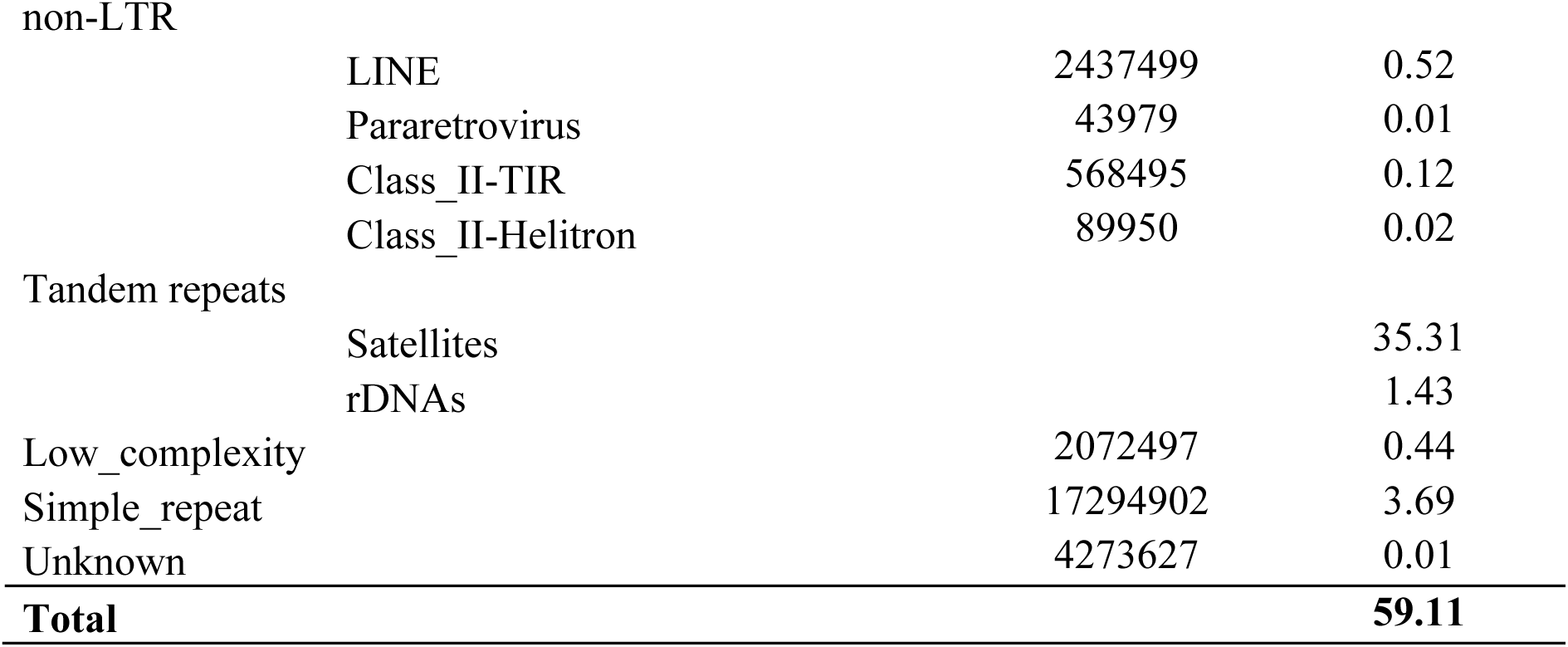
Genome proportion of the repetitive sequences in the genome assembly of *Luzula sylvatica* by DANTE-LTR and TAREAN.

| Element | Total length | Proportion |
| --- | --- | --- |
| LTR |  |  |
| Ty1-copia/Angela | 47526670 | 10.15 |
| Ty3-gypsy/non-chromovirus/Athila | 12206459 | 2.61 |
| Ty1-copia/SIRE | 7396917 | 1.58 |
| Ty1-copia/Ivana | 4937954 | 1.05 |
| Ty1-copia/Bianca | 3399411 | 0.73 |
| Ty1-copia/Ale | 1844668 | 0.39 |
| Ty1-copia/Tork | 1567483 | 0.33 |
| Ty3-gypsy/chromovirus/Reina | 1001818 | 0.21 |
| Ty3-gypsy/chromovirus/Tekay | 693833 | 0.15 |
| Ty1-copia/Ikeros | 689905 | 0.15 |
| Ty1-copia/TAR | 481869 | 0.10 |
| Ty1-copia/Alesia | 358026 | 0.08 |
| Ty3-gypsy/chromovirus/CRM | 176493 | 0.04 |

|  |  |  |  |
| --- | --- | --- | --- |
| non-LTR |  |  |  |
|  | LINE | 2437499 | 0.52 |
|  | Pararetrovirus | 43979 | 0.01 |
|  | Class_II-TIR | 568495 | 0.12 |
|  | Class_II-Helitron | 89950 | 0.02 |
| Tandem repeats |  |  |  |
|  | Satellites |  | 35.31 |
|  | rDNAs |  | 1.43 |
| Low_complexity |  | 2072497 | 0.44 |
| Simple_repeat |  | 17294902 | 3.69 |
| Unknown |  | 4273627 | 0.01 |
| **Total** |  |  | **59.11** |

Satellite DNA (satDNA) families’ distribution varied across the genome (**Fig. 2a**). *Lusy1* was spread throughout all pseudochromosomes, with higher densities in interstitial regions (**Fig. 2a**). In contrast, the other satDNAs (LsylSAT3–6) were found preferentially near telomeres or irregularly distributed on the chromosomes (**Fig. 2a and Table S2**). We localized *in situ* the two most abundant putative repeats (*Lusy1* and *Lusy2*) to corroborate the pattern obtained *in silico*. Remarkably, we observed that *Lusy1* shows a line-like distribution across the entire length of each sister chromatid, in a similar pattern to other holocentromeric repeats (Marques *et al*. 2015). In contrast, *Lusy2*, which showed a more irregular pattern, sometimes presented internal accumulations compared to *Lusy1* (**Fig. 2b**).

To investigate whether *L. sylvatica* holocentromeres are repeat-based, we performed a comparison analysis of these satellites with CENH3 ChIP-seq data. CENH3-ChIP-seq showed CENH3 enrichment for *Lusy1* and *Lusy2* repeats and depletion in LTR transposable elements throughout the *L. sylvatica* genome (**Fig. 2c**). Methylation was similar between *Lusy1/2* sequences, being highly enriched in CpG and CHG contexts at levels comparable to those of TEs. Regulatory sequences flanking the transcribed region of genes were sharply depleted of CpG methylation compared to intergenic regions and centers of the gene bodies (**Fig. 2c**). As recently reported for *Rhynchospora* (Hofstatter *et al*. 2022), CHG (but also CHH) methylation seems to increase toward the borders of *Lusy* satellites (**Fig. 2c**), reinforcing the idea of an evolutionary conserved epigenetic regulation of repeat-based holocentromeres in this lineage. By overlapping the annotation of the *Lusy1* and *Lusy2* centromeric repeats with CENH3 domains, we observed that centromeric units are mainly composed of *Lusy1* (n=232 out of 358 domains) and/or *Lusy2* sequences (n=96) (**Fig. 2d-e; Fig. S4**). Additionally, we found a small subset of CENH3 domains (n=33) associated with satellite-free regions, which were mainly composed of low-complexity repeats (53 %) and LTR-TEs (42.6 %) (**Fig. 2e; Fig. S5**). Although satellite-free CENH3 domains were depleted of genes, they often contain transposable elements (16 out of 33 domains; **Fig. 2e**). Remarkably, *Athila* elements belonging to *Ty3-gypsy* family were the most abundant, making up nearly 18 % of the length of the satellite-free CENH3 domains while representing only ∼3 % of the genome (**Table S3**). Satellite-free CENH3 domains were also positively correlated with CpG, CHG, and CHH methylation, similar to *Ty3-gypsy* TEs (**Fig. 2f; Fig. S6)**, suggesting that these two features occupy the same genomic niche. The presence of satellite-associated CENH3 domains was further confirmed by immuno-FISH analyses, where satellite *Lusy1* signals partially co-localize with CENH3 domains along the chromosome (**Fig. 3a-b**). Therefore, *L. sylvatica* represents a new case of repeat-based holocentromeres that are mostly, but not exclusively, composed of *Lusy1* repeats.

**Fig. 3:**
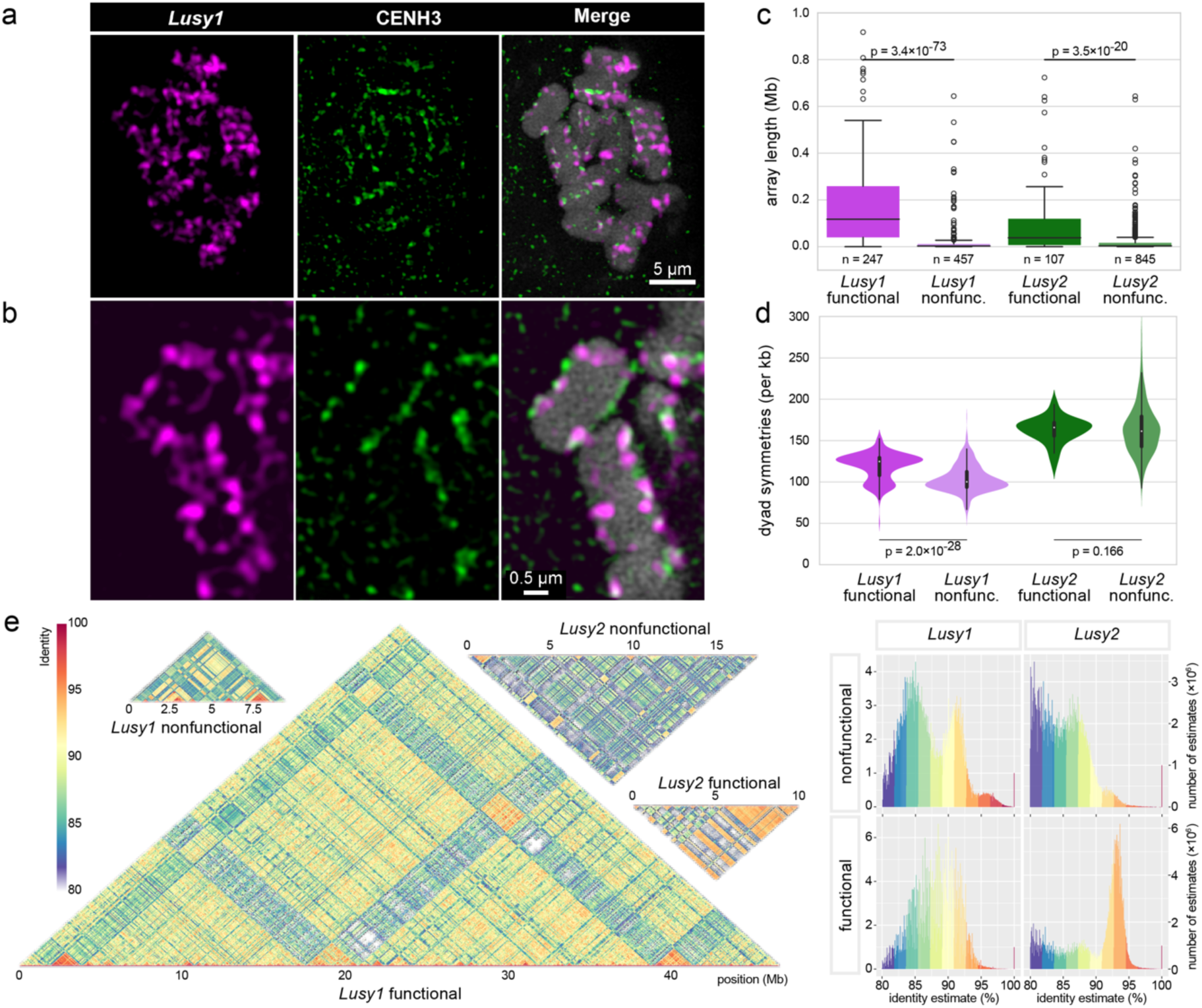
Association of CENH3 with centromeric satellite arrays. **(a-b)** Immuno-FISH showing partial colocalization of *Lusy1* repeats (magenta) and CENH3 (green) in metaphase chromosomes (counterstained with DAPI, blue). **(c)** Box plot of sizes of centromeric satellites *Lusy1* and *Lusy2* arrays associated with CENH3 (functional) or non-centromeric (nonfunctional). Statistical significance was tested using two-tailed Mann-Whitney U test. **(d)** Abundance of dyad symmetries in functional and nonfunctional arrays of centromeric satellites. Statistical significance was tested using one-tailed Mann-Whitney U test. Array counts are identical to subfigure **c**. **(e)** Homogeneity of functional array fragments (regions overlapping CENH3 domains) and whole nonfunctional arrays of centromeric satellites *Lusy1* and *Lusy2*. Dot plots showing sequence similarity between groups of concatenated arrays from the entire genome (left), histograms showing the frequency distribution of similarity values (right). Images in **(a)** and **(b)** represent single slices of 3D-SIM image stacks. Dot plots are shown proportional to their genomic abundance.

Using *in silico* mapping data, we also identified arrays of satellites *Lusy1* and *Lusy2* that lack association with CENH3 (hereafter referred to as nonfunctional). For *Lusy1*, 247 out of 704 arrays (35 %) are overlapping with CENH3 domains (functional). The length of these overlapping regions was 47 Mb out of 77 Mb in total (60 %). For *Lusy2*, 107 out of 952 arrays (11 %) contained CENH3 domains, making up 10 Mb out of 43 Mb (24 %) of total length. Nonfunctional arrays tended to be smaller than the functional arrays of the same satellite family, with an average length of functional/nonfunctional arrays of 189 kb/20 kb and 94 kb/20 kb for *Lusy1* and *Lusy2*, respectively (**Fig. 3c**). Functional *Lusy1* arrays contained a higher abundance of dyad symmetries. In *Lusy2,* this difference was not significant (**Fig. 3d**). Functional and nonfunctional arrays also differ in their inter-array sequence similarity. Functional arrays of both *Lusy1* and *Lusy2* satellites had higher average similarity across discrete arrays compared to nonfunctional arrays (88.0 vs 87.2 % and 89.6 vs 85.1 % for *Lusy1* and *Lusy2*, respectively; Fig. **3e**). Interestingly, nonfunctional *Lusy1* arrays show a clear bimodal distribution with one of the groups having a higher similarity than the corresponding functional arrays (**Fig. 3e**). Epigenetic status of the functional array chromatin also shows a striking contrast, since functional centromeric regions (i.e., *Lusy1* and *Lusy2* functional arrays, satellite-free centromeric units) are enriched with heterochromatin mark H3K9me2 and depleted of euchromatin mark H3K4me3, while the nonfunctional arrays are the opposite (**Fig. 2f and 6e**).

### Kinetochore proteins KNL1 and NDC80 remain conserved during the transition from mono-to holocentricity in *Luzula*

In *Cuscuta,* the transition to holocentricity was associated with massive changes in the localization of CENH3 and the kinetochore proteins KNL1, MIS12 and NDC80, representing the three complexes of the KMN network (Neumann *et al*. 2023; Oliveira *et al*. 2024). Unlike *Cuscuta* species, *Luzula* still possesses centromeric activity associated with CENH3 (**Fig. 1d and S7a**) (Nagaki *et al*. 2005; Heckmann *et al*. 2011). To test whether the kinetochore assembles along the poleward chromosome surface, as expected for holocentric chromosomes, we examined the localization of KNL1 and NDC80 in two holocentric woodrushes, *L. sylvatica* and *L. nivea* as well as in the related monocentric common rush *J. effusus*. Antibodies against KNL1 showed a similar pattern in both *L. sylvatica* and *L. nivea*, with signals detected as multiple clusters along the poleward surface of chromosomes, where microtubules attach (**Fig. 4a and S7a**). In addition, FISH signals from the centromeric repeat *Lusy1* were observed in the innermost part of KNL1 in a clustered pattern, contrasting with the continuous lines observed for KNL1, where microtubules attach. (**Fig. 4b)**. However, NDC80 signals were observed only in *L. nivea* (**Fig. S7a).** In *J. effusus*, KNL1 showed a specific dot-like localization in the primary constriction region of the chromosome, also associated with microtubule attachment sites (**Fig. S7b**). KNL1 and NDC80 provide spindle-binding sites detected by antibodies against α-tubulin, indicating that both proteins have a conserved kinetochore function in *Luzula* and *Juncus*.

**Fig. 4:**
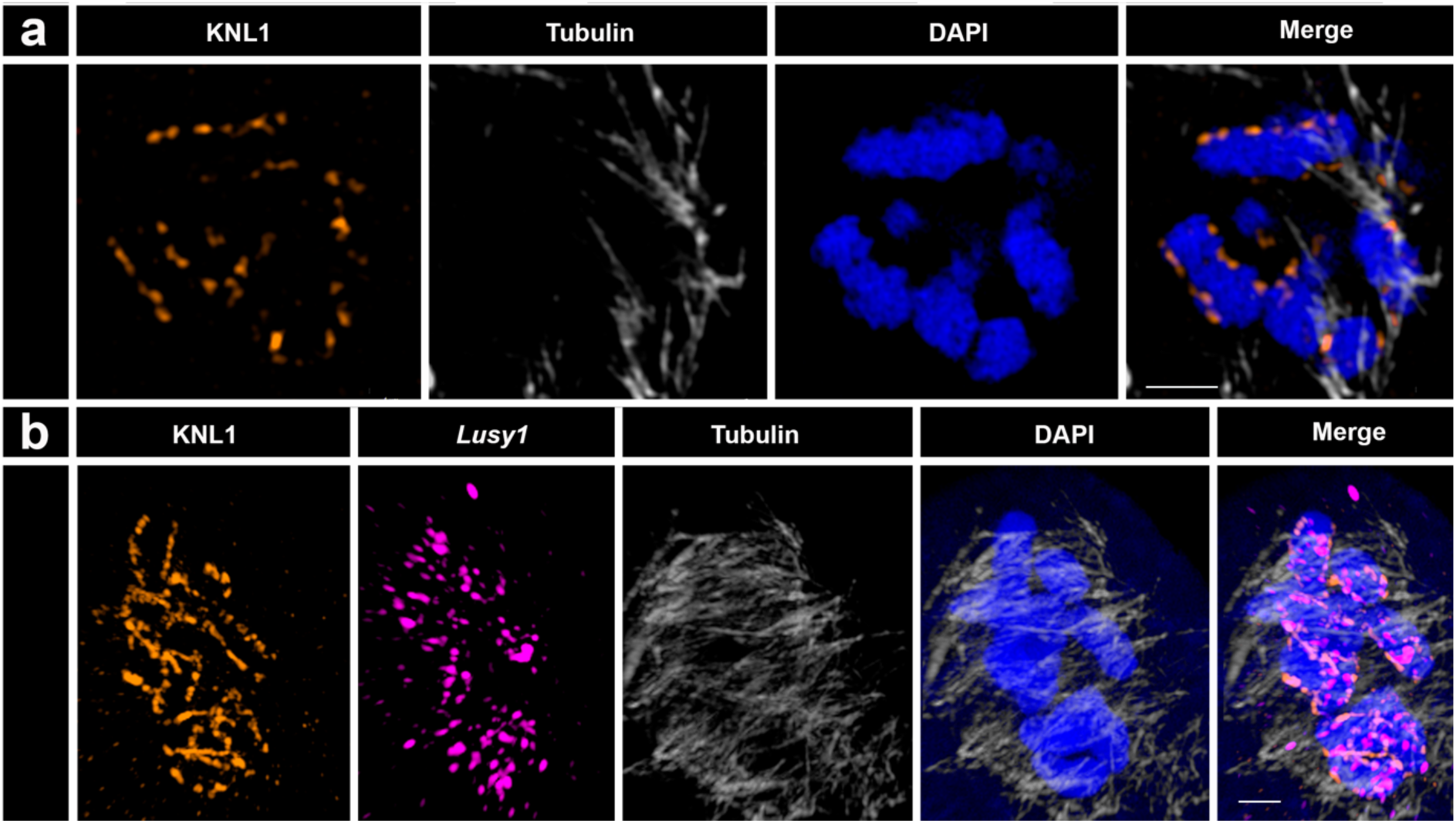
Detection of α-tubulin and kinetochore proteins in metaphase chromosomes of *L. sylvatica*. **(a)** KNL1 proteins localize specifically to the centromere surface, where microtubules (gray) bind. **(b)** Detection of KNL1, *Lusy1* repeats, and α-tubulin. Ends of microtubules colocalize with KNL1 proteins; some *Lusy1* sequences occur in the inner part of some kinetochore regions. Maximum intensity projections of 3D-SIM image stacks.

### Evolutionary dynamics of the centromeric satellites in *Luzula* species

To determine whether repeat-holocentromeres are conserved across other species of the genus *Luzula*, both an individual and comparative analysis of the repeatome using RepeatExplorer was performed in 13 species, including *L. sylvatica* (**Table S4-S6**). The global genomic proportion of repetitive DNA varied from 35.24 % (*L. pilosa*) to 66.29 % (*L. wahlenbergii*) (**Table S5**). In general, satDNAs were the most abundant class of repeats, comprising up to 49 % of the *L. sudetica* genome. The centromeric satellite *Lusy1* was one of the most abundant among all satellites, representing up to 47.41 % of *L. sudetica* genome but only 3.33 % of the *L. nivea* genome (**Table S5**). *Lusy2* also showed variation in abundance among species, ranging from 0.34 % (*L. multiflora* subsp. *frigida*) to 31.03 % (*L. luzuloides*), being also found in *L. elegans* genome (0.72 %; **Table S5**), a species with previously undetected holocentromeric repeat (Heckmann et al. 2013). LTR retrotransposons revealed variable abundances among species, with the *Ty1-copia* superfamily being the most represented (1.21 % in *L. pilosa* to 41.35 % in *L. elegans*; **Table S5**).

Comparative repeat analysis resulted in 166 shared clusters (**Fig. 5a, Fig. S8, Table S6**). Variants of *Lusy1,* the most abundant satellite family in *Luzula*, were found in all analyzed species, except in *L. elegans*, where this satellite was not detected even in an additional fine search of the raw sequencing reads (**Fig. 5a**). Different variants of the *Ty1-copia Angela* lineage were found in high abundance among the species, being more dominant in the genomes of *L. arcuata* and *L. elegans* (**Fig. 5a; Table S6**). *Ivana* and *SIRE Ty1-copia* lineages were also shared among all species, although they exhibited lower abundance than *Angela* (**Table S5-6**).

**Fig. 5:**
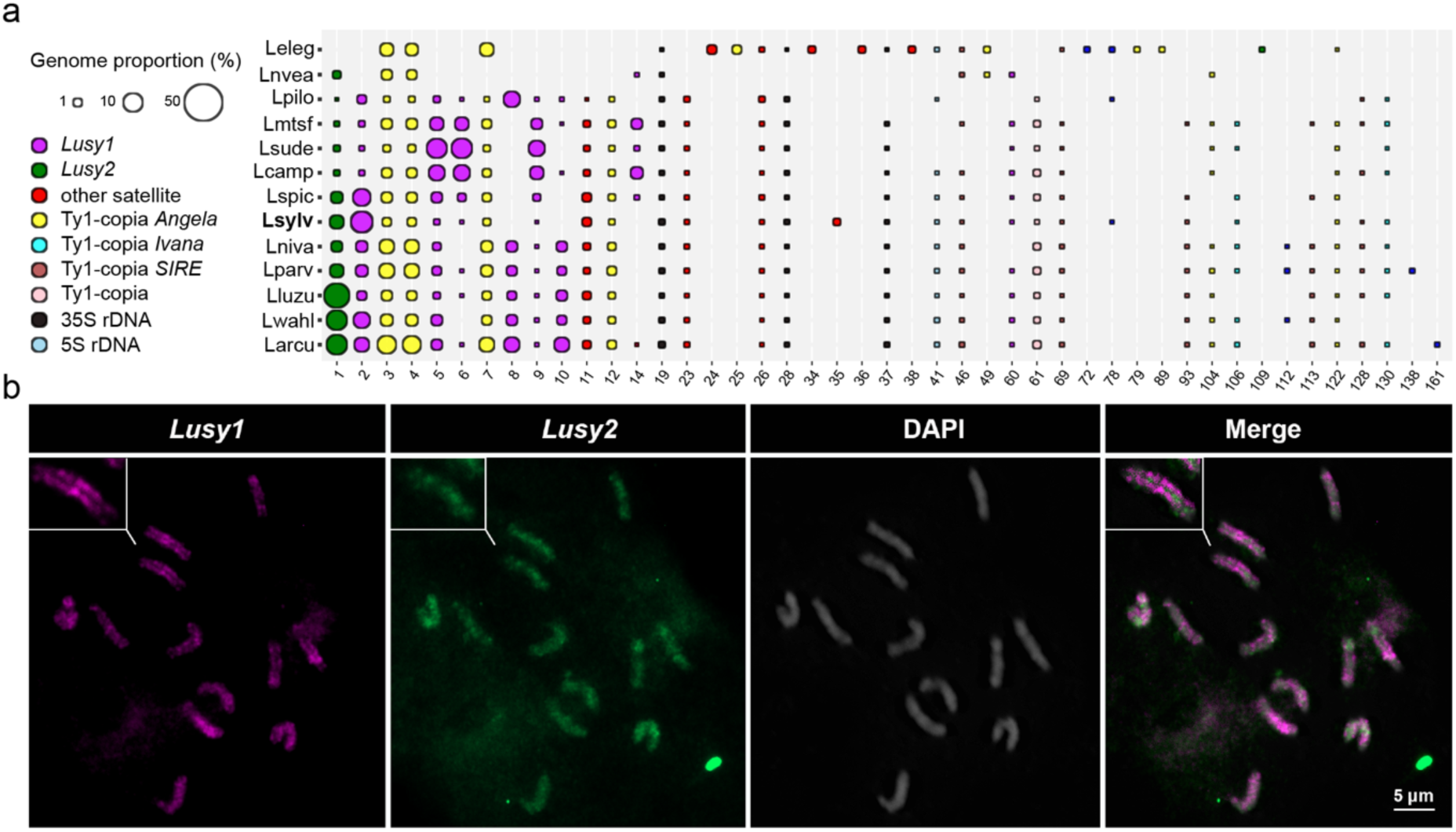
Comparative analyses of the main types of repetitive sequences in *Luzula* species. **(a)** Comparative analyses of the abundance of the main types of repetitive sequences in *Luzula* species. Code names correspond to Larcu: *Luzula arcuata*, Lcamp: *Luzula campestris*, Leleg: *Luzula elegans*, Lluzu: *Luzula luzuloides*, Lmtsf: *Luzula multiflora* subsp. *frigida*, Lniva: *Luzula nivalis*, Lnvea: *Luzula nivea*, Lparv: *Luzula parviflora*, Lpilo: *Luzula pilosa*, Lspic: *Luzula spicata*, Lsude: *Luzula sudetica*, Lsylv: *Luzula sylvatica*, Lwahl: *Luzula wahlenbergii*. The size of the ball is proportional to the genome abundance of that cluster for each species. The colors of the balls correspond to different repetitive sequence types (see Table S4 for details). **(b)** FISH showing a wide-spreading localization of satellite *Lusy1* (magenta) and a dispersed pattern of *Lusy2* (green) repeats in metaphase chromosomes of *L. nivea*. Chromosomes were counterstained with DAPI. Scale bar, 2 µm.

Because *Lusy1* and *Lusy2* were the most abundant satellites in the comparative analysis, consistent with the observation from the *L. sylvatica* genome, we performed FISH to confirm their distribution also in *L. nivea*, the species with the lowest abundance of *Lusy1*. Like *L. sylvatica*, the FISH signals of *Lusy1* in *L. nivea* showed a line-like distribution along the chromosomes. However, exhibiting both enriched and depleted labeled chromosomal regions. Furthermore, *Lusy2* showed clustered signals enriched at interstitial and terminal regions in a non-linear pattern (**Fig. 5b**). These results suggest a similar repeat-based holocentromere organization for other *Luzula* species as well.

### Chromosome fusions drive karyotypic evolution in *Luzula*

Chromosomes from some grasses and several holocentric species have undergone extensive karyotypic rearrangements through fusions (Wang et al., 2021; Hofstatter *et al*., 2022; Escudero et al. 2023). To investigate the possible association between holocentricity and chromosome fusions, we analyzed synteny between the genomes of the holocentric *L. sylvatica* and the monocentric *J. effusus* species (**Fig. 6, Fig. S9**). Considering *n* = 20 as the putative ancestral karyotype for the family Juncaceae (Drábková 2013), the synteny analysis between the two genomes revealed that the chromosomes of *L. sylvatica* consist of fused blocks from *J. effusus* chromosomes (dysploid with *n* = 21), resulting in a descending dysploidy to *n* = 6 (**Fig. 6a**). Despite their high chromosome number and centromere-type differences, small arrangements and large syntenic blocks were identified between both genomes, indicating well-conserved genomic structures in this family, with a total of 86.4 % (23,016 gene pairs) of the *J. effusus* genome being syntenic with *L. sylvatica* (**Fig. 6b, Table S7, Fig. S9-10**).

**Fig. 6:**
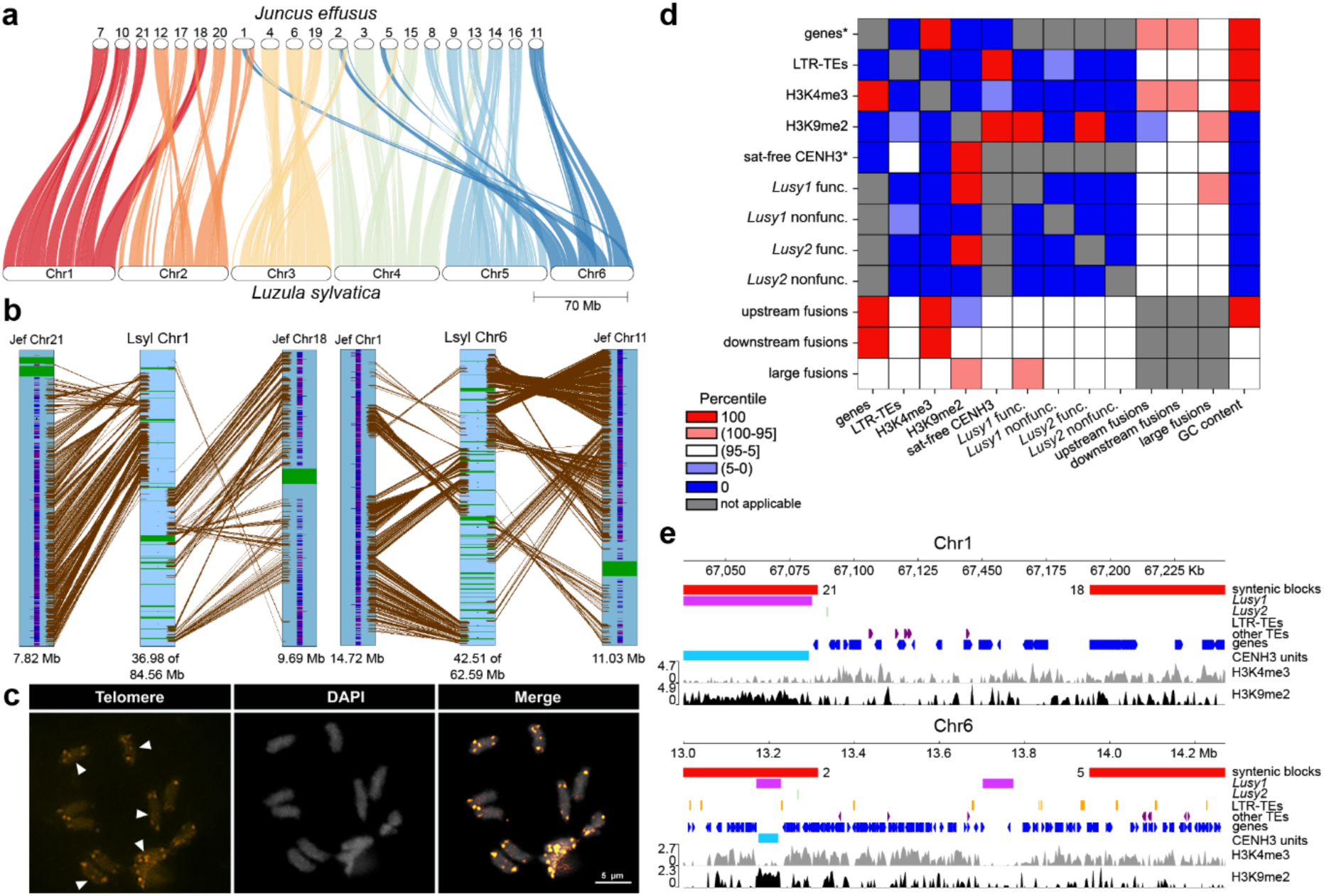
Genome synteny comparison of *L. sylvatica* and *J. effusus*. **(a)** Genome synteny patterns showing macro-conserved blocks of *J. effusus* that were fused into *Luzula* chromosomes. **(b)** *L. sylvatica* chromosome 1 and 6 (Lsyl) showing the fusion of syntenic blocks of *J. effusus* (Jef) 1, 11, 18 and 21. Genes, CENH3 and telomere domains are annotated as blue-purple, green and black stripes, respectively. **(c)** FISH of telomeric DNA showing chromosomes with remnant interstitial telomeric sites, suggesting ancestral fusion events. **(d)** Colocalization between major genomic and epigenomic features and fusion points (columns) based on comparison of overlap with simulated regions (rows). Heatmap values show the rank of real overlap values among a distribution of overlaps with simulated regions. * Signifies exclusion of satellite array positions from permitted simulated region locations, for the rest of the features, the whole genome was used for simulated regions. **(e)** Examples of fusion sites on Chr1 and Chr6 in *L. sylvatica*.

We have recently shown that the holocentromeric repeat *Tyba* can be involved in facilitating end-to-end chromosome fusions in *Rhynchospora* species (Hofstatter *et al*. 2022). To assess the possible role of *Lusy1* and *Lusy2* in the fusion points observed in *Luzula* genomes, we looked for specific enrichment of these repeats and telomeric repeats at the fusion points. We found evidence for interstitial telomeric sites (ITS) using FISH (**Fig. 6c**), while the inspection of the pseudomolecules revealed three ITS in total, in Chr1, Chr2 and Chr5. However, in none of these cases, the ITS site was correlated with a synteny block transition between different *Juncus* chromosomes (**Fig. S11**). Further, the size of the fusion points in *L. sylvatica*, defined as the space between syntenic blocks, revealed a size range from 10 kb to 8 Mb. At the block boundaries (flanking regions of 50 kb), we observed a positive association with genes (**Fig. 6d, S12**). Large fusion points (>100 kb) also contained satellites and/or transposable elements (**Fig. 6e**). However, the enrichment of the large fusion points with repeats is not prominent enough to be recognized at the scale of genome-wide colocalization between features (except possible enrichment for functional *Lusy1* arrays; **Fig. 6d**).

## Discussion

Holocentromeres have evolved from a monocentric ancestor multiple times during the evolution of eukaryotes, and despite the convergent appearance of extended centromere, each of these events results in specific genome organization and adaptation of the kinetochore protein machinery (Márquez-Corro *et al*. 2019; Senaratne *et al*. 2022; Hofstatter *et al*. 2022; Lucek *et al*. 2022; Kuo *et al*. 2023). In *Cuscuta*, holocentricity co-occurs with CENH3-independent mitotic spindle attachment and extensive changes in kinetochore structural and regulatory protein genes (Oliveira *et al*. 2020; Neumann *et al*. 2023). In insects, transitions to holocentricity are associated with the loss of CENH3 domains while the inner kinetochore complex remains relatively conserved (Drinnenberg *et al*. 2014; Cortes-Silva *et al*. 2020). In cases where the kinetochore protein localization and function are maintained, their localization can present a spectrum between continuous line-like and discrete cluster-like distribution along the chromosome (Schubert *et al*. 2020). While previously studied *Luzula* species (*L. elegans* and *L. nivea*) display a line-like distribution of CENH3 (Nagaki *et al*. 2005; Heckmann *et al*. 2011; Oliveira *et al*. 2024), we detected signals more resembling cluster-like distribution for CENH3 and KNL1 proteins in *L. sylvatica*, suggesting a more discontinuous holocentromere organization in this species.

We show that functional holocentromeres in *L. sylvatica* are mainly made of *Lusy1* and partially *Lusy2* satellite repeats organized as kilobase-scale, non-uniformly spaced CENH3-positive centromeric units. However, the presence of several repeat-less centromere units suggests a more complex determination of centromere function in this species that needs to be further examined. These units display heterochromatin epigenetic characteristics and alternate with euchromatin, rich with coding regions, along the chromosome. Previous studies have identified a potential 178 bp centromeric tandem repeat in *L. nivea* and other *Luzula* species with similarity to the centromeric satellite *RCS2* from rice (Haizel *et al*. 2005). Indeed, the 178 bp satellite shares 87% similarity with the *Lusy2* satellite, which we found in all species. Although *Lusy2* shows a partial enrichment in centromeric regions, it does not cover the entirety of the chromosomes unlike 124 bp *Lusy1*, which largely encompasses the entire holocentromere of *L. sylvatica* and is abundant in most analyzed *Luzula* species, suggesting that *Lusy1* is the primary centromeric satellite of the genus. A notable exception is an early-diverging species *L. elegans*, where *Lusy1* is absent and *Lusy2* represents <1 % of the genome. Remarkably, none of the 20 previously analyzed satellite repeats in *L. elegans* exhibited a centromeric pattern, despite repetitive sequences making up ∼60 % of the genome (Heckmann *et al*. 2013; Jankowska *et al*. 2015). Repeat-based holocentromeres have been previously reported in *Chionographis japonica* (Kuo *et al*. 2023), *Rhynchospora* (Marques *et al*. 2015; Hofstatter *et al*. 2022; Castellani *et al*. 2024), in the nematode *Meloidogyne incognita* (Despot-Slade *et al*. 2021), and in mulberry (Ma *et al*. 2023). Remarkably, previous studies did not detect the holocentromeric repeat *Tyba* in early diverging lineages in *Rhynchospora*, and instead have diverse satellite arrays arranged in block-like patterns (Ribeiro *et al*. 2016; Costa *et al*. 2021), which mirrors the lack of *Lusy1* in *L. elegans*. In both of these genera, the colonization of holocentromeres by contemporary genus-specific centromeric satellite families occurred only after the process of transition to holocentricity began (Costa *et al*. 2023).

This raises questions about the expansion and functional role of satellites in holocentromeres. A process similar to the establishment of neocentromeres in monocentrics could be taking place. Neocentromeres can arise in heterogeneous genomic regions that become subject to rapid cycles of invasion and purification of repetitive sequences through satellite homogenization (Wlodzimierz *et al*. 2023). In *L. sylvatica*, we have identified several satellite-free centromeric units reminiscent of maize *de novo* centromeres in their gene-poor region targeting, CHG and CHH methylation, and possible association with *Ty3-gypsy* elements. These units could then become a subject of competition for centromere dominance between *Lusy* satellites driven by satellite homogenization and evolutionary selection pressure for centromere stabilizing effects such as advantages in CENH3 loading (Kasinathan and Henikoff 2018) or nucleosome formation and positioning (Talbert and Henikoff 2020; Hofstatter *et al*. 2022; Costa *et al*. 2023). An analogous process of acquisition of heterochromatin epigenetic modifications, accumulation of transposable elements, and invasion of satellite repeats has been described in monocentrics as evolutionary new centromere maturation (Piras *et al*. 2010; Cappelletti *et al*. 2022).

Furthermore, polycentric chromosomes can also originate from monocentromere spreading or chromosome rearrangements. The transition to holocentricity in *Luzula* could have been initiated earlier than the split of the *Luzula*/*Juncus* genus, since the repeat-based centromeres of *J. effusus* were recently described as an atypical monocentromere, with up to three centromere cores and different types of centromeric organization, resembling a wide-spread metapolycentric organization (Dias *et al*. 2024). In *Luzula*, dramatic karyotype changes took place, resulting in reduced chromosome number. Although phylogenetic relationships are poorly resolved in the genus, descending dysploidy has been observed in species from different *Luzula* clades (*L. elegans* in the Marlenia clade and *L. purpureo-splendens* in the Nodulosae clade), indicating that this process of chromosomal evolution has occurred independently at least twice during the evolution of the genus (Bozek *et al*. 2012). Our results support dysploidy as the main driver of karyotype evolution in holocentric organisms (**Fig. S10**; (Bozek *et al*. 2012; Guerra 2016; Senaratne *et al*. 2022), since fission and fusion events have been described in sedges, leading to dysploid karyotypes in the holocentric genera *Rhynchospora* (Hofstatter *et al*. 2022; Mata-Sucre *et al*. 2024) and *Carex* (Ning *et al*. 2023; Escudero *et al*. 2023), as well in some holocentric butterflies (Cicconardi *et al*. 2021; Wright *et al*. 2024). However, the fusion of ancestral chromosomes that resemble chromosomes from the sister genus *Juncus* resulting in the dysploid *L. sylvatica* is intriguing, since it involves a simultaneous shift of centromere organization. The 21 putative *Juncus* ancestral-like chromosomes merged into six *L. sylvatica* chromosomes while undergoing additional chromosome rearrangements, genomic reshuffling, and repetitive DNA turnover in the past 56 million years of divergence (Elliott and Davies 2019). Unlike the repeat-mediated chromosome fusions observed in *Rhynchospora* (Hofstatter *et al*. 2022), fusion sites in *L. sylvatica* lack genomic footprints, suggesting another mechanism of chromosome fusions during karyotype evolution in *Luzula* and/or a masking over time by additional chromosomal rearrangements and dynamic repeat turnover.

Based on these findings, we propose a multistep model of evolutionary transition to holocentricity in the genus *Luzula* (**Fig. 7**). At first, a hypothesized change of the genomic processes responsible for centromere maintenance enables centromere spreading without inactivation, as observed in *Juncus*. Next, stable polycentric chromosomes arise through multiple chromosome fusions of *Juncus*-like chromosomes. Then, polycentric organization gradually transitions to line-like holocentric organization with tens to hundreds of discrete CENH3 domains through genome rearrangements and CENH3 spreading or seeding with subsequent centromere maturation, culminating with satellites invading these loci by a combination of satellite DNA library diversification and concerted evolution, as observed for *Tyba* repeats (Costa *et al*. 2023). From this point of view, the presence of satellite-free centromeric units, the uneven distribution of centromeric satellite repeats and the cluster-like distribution of CENH3 and outer kinetochore proteins in *L. sylvatica* can be interpreted as intermediate stages of ongoing holocentromere transition. Further research can provide insight into the molecular mechanisms’ adaptation taking place during the holocentric transition and its triggers.

**Fig. 7:**
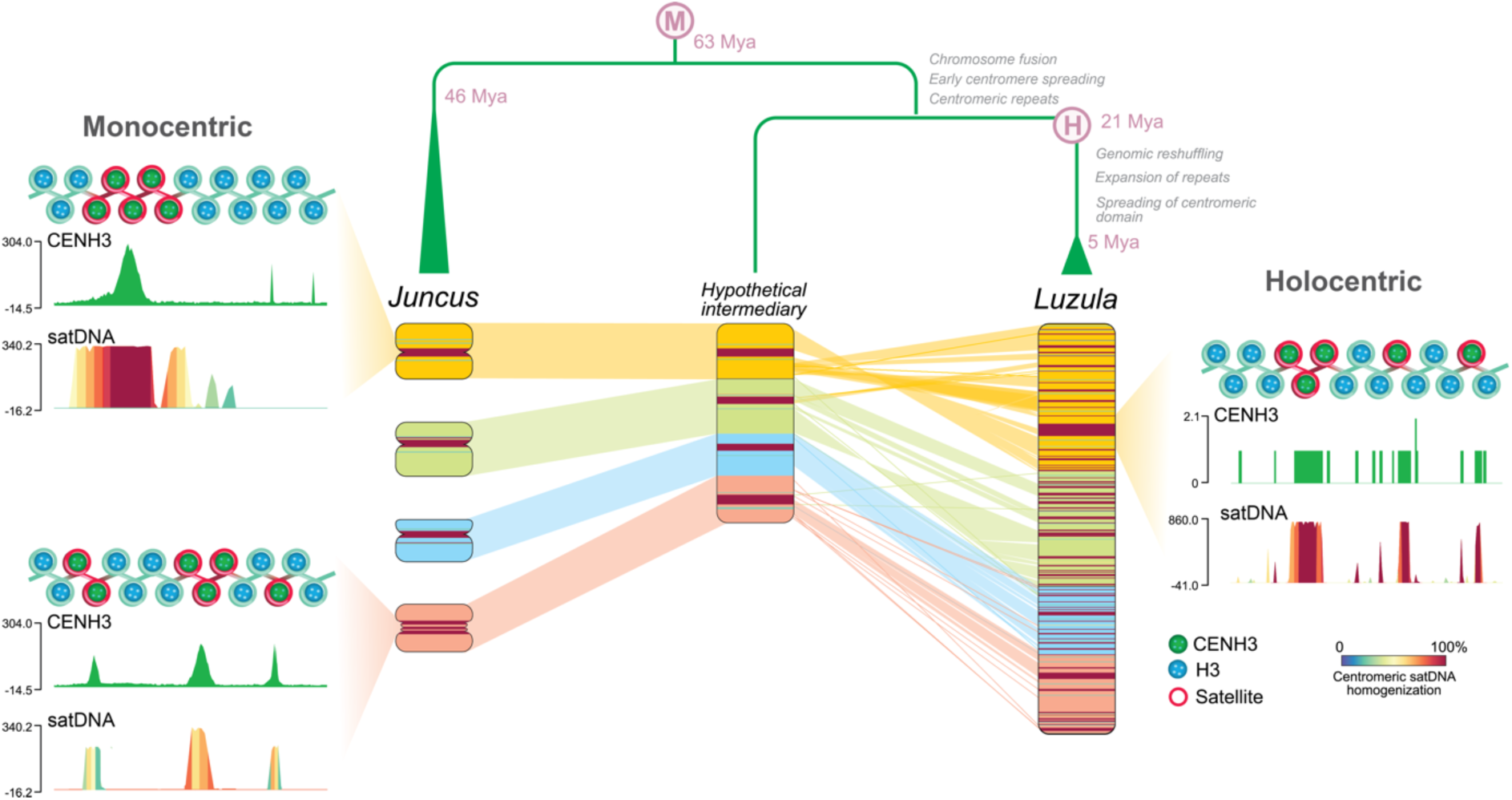
Model for the origin of holocentricity in woodrushes. After several fusions of whole atypical monocentric chromosomes (*Juncus-*like type), centromeric domains were initially conserved in the larger chromosomes (hypothetical intermediate state), forming polycentric chromosomes. Subsequently, expansion of the centromeric domain and genome rearrangement gave rise to the holocentric condition. Later colonization of *Lusy*-type satellites allowed the maintenance of functional centromeres. M: monocentric. H: holocentric. Divergence time was obtained from the Timetree of Life (https://timetree.org/).

## Acknowledgments

We thank Dra. Magdalena Vaio for her comments, which led to an improvement in the quality of the work. We acknowledge the excellent technical assistance of Ursula Pfordt, Laís Marinho and Christina Philipp. This work was funded by the Max Planck Society (core funding to A.M.), by the Deutsche Forschungsgemeinschaft (DFG, grant no. MA 9363/3-1 to A.M.) and by the European Union (European Research Council Starting Grant, HoloRECOMB, grant no. 101114879 to A.M.). DFG founded this work under Germany’s Excellence Strategy—EXC 2048/1–390686111 (A.M.). We thank the International Cooperation Program PROBRAL (CAPES/DAAD project number 88881.144086/2017-01) for the scholarship offered to Y.M.S. We thank the Darwin Tree of Life Project at the Wellcome Sanger Institute for making the data available at https://www.darwintreeoflife.org/project-resources. Computational resources for RepeatExplorer analysis were provided by the ELIXIR-CZ project (LM2023055), part of the international ELIXIR infrastructure. e-INFRA CZ project (ID:90254), supported by the Ministry of Education, Youth and Sports of the Czech Republic provided computational resources for the analysis of ChIP-seq data. The work of M.K. was supported by grant 21– 00580S from the Czech Science Foundation awarded to E.K. L.O. was supported by grand 20-25440S from Czech Science Foundation). A.P.H. and G.S. received productivity fellowship from CNPq (process numbers PQ-312852/2021-5 and PQ-312852/2021-5, respectively).

## Contributions

Y.M.S: Investigation, Validation, Formal analysis, Data Curation, Writing-Original draft preparation. M.K.: Validation, Formal analysis, Data Curation, Writing-Reviewing and Editing. L.O.: Investigation, Resources. P.N. and J.M.: Resources, Writing-Reviewing and Editing. V.S. and A.H.: Resources, Writing-Reviewing and Editing. E.K, A.P.H and G.S.: Supervision, Resources, Writing-Reviewing and Editing. A.M.: Conceptualization, Supervision, Resources, Funding acquisition, Writing-Reviewing and Editing.

## Data availability

All sequencing data used in this study have been deposited at NCBI under the BioProject ID XXXXXXX and are publicly available as of the date of publication. The reference genomes, sequencing data, annotations and all tracks presented in this work are made available for download at DRYAD: LINK. The REXdb database Viridiplantae v.3.0 [http://repeatexplorer.org] is publicly available. All other data needed to evaluate the conclusions in the paper are provided in the paper and/or the supplemental information.

## Code availability

The original code used in this study is available on GitHub at https://github.com/437364/Repeat-based-holocentromeres-of-Luzula-sylvatica. Any additional information required to re-analyze the data reported in this paper is available from the corresponding author upon request.

## Methods

### Plant material

For cytogenetic analyses, plants from natural populations of *L. sylvatica* were collected in Cologne, Germany, and further cultivated under controlled greenhouse conditions (16h daylight, 26 °C, >70% humidity). The ornamental plant *L. nivea* was commercially obtained (Dingers Gartencenter) and cultivated under controlled greenhouse conditions (16h daylight, 20°C).

### Genome assembly and Hi-C scaffolding

HiFi and Hi-C reads obtained through the Darwin Tree of Life database (www.darwintreeoflife.org) were assembled using Hifiasm (Cheng *et al*. 2021), available at https://github.com/chhylp123/hifiasm, following the command: “*hifiasm-o output.asm-t 40 reads.fq.gz*”. Preliminary assemblies were evaluated for contiguity and completeness with BUSCO (Seppey *et al*. 2019) and QUAST (Gurevich *et al*. 2013).

Hi-C reads were first mapped to the primary contigs file obtained from the Hifiasm assembler using BWA (Li and Durbin 2009) following the hic-pipeline (https://github.com/esrice/hic-pipeline). Hi-C scaffolding was performed using SALSA2 (https://github.com/marbl/SALSA) (Ghurye *et al*. 2019) with default parameters using ‘*GATC, GAATC, GATTC, GAGTC, GACTC*’ as restriction sites. After testing several minimum mapping quality values of bam alignments, the final scaffolding was performed with MAPQ10. Several rounds of assembly correction guided by Hi-C contact maps and manual curation of scaffolds were performed to obtain the six pseudomolecules.

Genome size estimate was obtained from HiFi reads using findGSE (Sun *et al*. 2018). First, a histogram of k-mers was created using jellyfish (Marçais and Kingsford 2011), and then the findGSE R package was used for model fitting according to package documentation (https://github.com/schneebergerlab/findGSE).

### Chromatin immunoprecipitation (ChIPseq) sequencing and analysis

ChIP experiments were performed following Hofstatter *et al*. (2022). *L. sylvatica* leaves were harvested and frozen in liquid nitrogen until sufficient material was obtained. The samples were fixed in 4% formaldehyde for 30 min and the chromatin was sonicated to enrich for 300 bp fragments. Then, 40 ng of sonicated chromatin was incubated with 2 ng of antibody overnight. Immunoprecipitation experiments were carried out for the rabbit anti-*L. elegans* CENH3 (LeCENH3; Ma et al., 2016), rabbit anti-H3K4me3 (abcam, ab8580), and mouse anti-H3K9me2 (abcam, ab1220). Anti-LeCENH3 that was originally developed against 3-RTKHFSNRKSIPPKKQTPAK-23 peptide from *Luzula elegans* bears 65% similarity to the corresponding sequence 3-RTKHFSLRSRHPKKQRTAA-22 from *Luzula sylvatica* CENH3 (GenBank: KJ934236.1). Recombinant rabbit IgG (abcam, ab172730) and no-antibody inputs were used as controls. Two experimental replications were also maintained for all the combinations. ChIP DNA was quality-controlled using the NGS-assay on a FEMTO-pulse (Agilent); next, an Illumina-compatible library was prepared for all immunoprecipitants with the Ovation Ultralow V2 DNA-Seq library preparation kit (Tecan Genomics) and single-end 1 x 150-bp reads were sequenced on a NextSeq 2000 (Illumina) device. For each library, an average of 20 million reads was obtained.

The raw sequencing reads were trimmed by Cutadapt (Martin 2011) to remove low-quality nucleotides (with quality score less than 20) and adapters. Trimmed ChIPed 150-bp single-end reads were mapped to the respective reference genome with bowtie2 (Langmead and Salzberg 2012), where all read duplicates were removed and only the single best matching read was kept on the final alignment BAM file. ChIP vs input signal was calculated as the log2 ratio of read coverages normalized by reads per kilobase per million mapped reads (RPKM) using the bamCompare tool from deepTools package (Ramírez *et al*. 2016). Averaged signal from both replicates was visualized using pyGenomeTracks (Lopez-Delisle *et al*. 2021). Metaplots obtained by the plotProfile function from deepTools were used to compare the distribution with other genomic features (Ramírez *et al*. 2016). To concretize enriched domains, we performed peak-calling by MACS3 (Zhang *et al*. 2008) and epic2 (Stovner and Sætrom 2019) and filtered only the peaks identified by both tools in both replicates. This high stringency peak filtering approach was chosen to reduce the risk of including false positive CENH3 domains in subsequent analyses. Based on analysis of CENH3 peak clustering, peaks closer than 150 kb were merged to obtain uninterrupted centromeric units (code available on GitHub at https://github.com/437364/Repeat-based-holocentromeres-of-Luzula-sylvatica).

### Methylation sequencing and analysis

To analyze DNA methylation level, a sequencing library was prepared using NEBNext® Enzymatic Methyl-seq Kit (NEB; catalog number E7120S). Library sequencing was performed using NextSeq 2000 (Illumina) platform, obtaining ∼20 M paired-end reads. Sequencing data was analyzed using the Bismarck pipeline (Krueger and Andrews 2011) according to the toolkit documentation (https://felixkrueger.github.io/Bismark/bismark/). Coverage files for CpG, CHG, and CHH methylation contexts were converted to bigwig.

### Synteny analysis

The synteny analysis between *L. sylvatica* and *J. effusus* was performed with CoGe SynMap platform (https://genomevolution.org/coge/SynMap.pl) (Lyons *et al*. 2008) and SyMAP v. 5.0.6 (Soderlund *et al*. 2006, 2011). For this analysis, CDS sequences, centromeric and telomeric repeats of both species were used. Synteny plots were obtained with GENESPACE (https://github.com/jtlovell/GENESPACE) (Lovell *et al*. 2022). Orthologs were identified following the steps: (1) using the BlastZ tool; (2) synteny analysis was performed using DAGChainer, using 25 genes as the maximum distance between two matches (-D) and 20 genes as the minimum number of aligned pairs (-A); (3) Quota Align Merge was used to merge syntenic blocks, with 50 genes as the maximum distance between them; and (4) orthologous and paralogous blocks were differentiated according to the synonymous substitution rate (Ks) using CodeML (where 2 was the maximum value of log10), and represented with different colors in the dot plot (**Fig. S7a**).

For the characterization of the regions involved in fusion points, we followed (Hofstatter *et al*. 2022). The synteny alignment between *L. sylvatica* and *J. effusus* genomes obtained in SyMAP allowed us to pin the putative regions around the borders of the fusion events. In order, to identify the underlying sequences at the fusion regions, we loaded annotation features for genes, TEs, and tandem repeats on SyMAP alignments. This allowed us to detect the sequence types in the putative fused regions. Further inspection and characterization of such regions were done by checking the genome coordinates and annotation features with Geneious (Kearse *et al*. 2012).

To further analyze colocalization of genomic features, epigenetic marks, and fusion points; we selected 50kb regions upstream and downstream of syntenic block edges facing the fusion point and also fusion points where the space between two syntenic blocks was larger than 100 kb. To analyze whether these regions are enriched or depleted of specific features, 1000 rounds of random region distribution or random region distribution excluding satellite array locations were simulated (this was done to improve the reliability of the null distribution for features that are defined as not overlapping with satellite arrays, e.g. satellite-free CENH3 domains and genes) (Kanduri *et al*. 2019) using *bedtools shuffle*. Then, overlap with all other studied features was calculated for simulated and real regions as a proportion of overlapping bases to all bases covered by the feature. The percentage of real overlap proportion in the distribution of simulated values was reported (code available on GitHub at https://github.com/437364/Repeat-based-holocentromeres-of-Luzula-sylvatica).

### Repeat characterization

Available Illumina reads from ENA (<u>ENA Browser (ebi.ac.uk)</u>) was filtered by quality with 95 % of bases equal to or above the quality cut-off value of 10 using RepeatExplorer2 pipeline (https://repeatexplorer-elixir.cerit-sc.cz/) (Novák *et al*. 2020). The clustering was performed using the default settings of 90 % similarity over 55 % of the read length. For the comparative analyses, we performed an all-to-all similarity comparison across all species following the same approach. Because the genome size is unknown for some analyzed species, each set of reads was down sampled to 1,000,000 for each species. Additionally, a subsample of eight species with known genome size were analyzed to compare the results. Samples from each species were identified with the four-letter prefixes shown in Table 1, and concatenated to produce datasets as input for RepeatExplorer2 (Novák *et al*. 2020) graph-based clustering.

The automatic annotation of repeat clusters was manually inspected and revised, and followed by a recalculation of the genome proportion of each repeat type where applicable. DANTE and DANTE-LTR annotation pipeline was used to identify full-length LTR retrotransposons in the assembled genome, containing a set of protein domains from REXdb (Novák *et al*. 2020). Overall repeat composition was calculated excluding clusters of organelle DNA (chloroplast and mitochondrial DNA). All contigs with tandem repetitions identified by TAREAN (Novák *et al*. 2017), were compared to verify their homology with DOTTER (Sonnhammer and Durbin 1995). All repeat features were individually mapped to the genome by BLAST using Geneious (Kearse *et al*. 2012), converted to BED and used as an input track for a genome-wide overview with ShinyCircos (Yu *et al*. 2018). Interstitial telomere sequences (ITS) were annotated by searching for regions longer than 100 bp with at least 90% similarity to consensus *Arabidopsis*-type telomere (TTTAGGG).

### Characterization of centromeric units

Centromeric units from ChIP-seq analysis were grouped by chromosome and their size, count, and density were calculated. Next, centromeric units were overlapped with locations of satellites to obtain locations and extract sequences of functional array fragments (precise regions where centromeric satellites *Lusy1/Lusy2* and CENH3 domains overlap), nonfunctional arrays (whole *Lusy1/Lusy2* arrays not overlapping CENH3 domains), and satellite-free units (CENH3 domains not overlapping any satellites). Sequences of discrete arrays of each type were concatenated and their homogeneity assessed using ModDotPlot (https://github.com/marbl/ModDotPlot). Dyad symmetries were identified in each array using the EMBOSS palindrome tool as described in (Kasinathan and Henikoff 2018) with *-nummismatches parameter set to 0.* Statistical significance of the increase of dyad symmetry abundance for functional arrays was tested using one-tailed Mann-Whitney U test from scipy package (Virtanen et al., 2020). Proportions of functional and nonfunctional arrays as well as other genetic and epigenetic features in 100kb windows were correlated using Spearman’s rank correlation from scipy package and resulting correlation coefficients were plotted in a heatmap (code available on GitHub at https://github.com/437364/Repeat-based-holocentromeres-of-Luzula-sylvatica). Additional packages were used for data handling and visualization (Hunter 2007; McKinney and others 2011; Dale *et al*. 2011; Harris *et al*. 2020; Waskom 2021).

### Cytogenetic and immunostaining of CENH3 protein

Mitotic preparations were made from root meristems fixed in 4% paraformaldehyde and Tris buffer (10 mM Tris, 10 mM EDTA, 100 mM NaCl, 0.1% Triton, pH 7.5) for 30 min on ice in vacuum and for another 20 min only on ice. After washing twice in 1 x PBS for 10 min, the roots were digested in a cellulase-pectinase (2% w/v /20% v/v solution) containing PBS buffer and squashed in PBS. The coverslips were removed in liquid nitrogen and the slides were air dried and stained in 2 µg/mL DAPI/Vectashield mounting medium for slide selection under the epifluorescence microscope. The slides with the highest number of cells in division were incubated in 3% (w/v) bovine serum albumin (BSA) containing 0.1% Triton X-100 in PBS. Immunostaining was performed using the primary antibodies rabbit anti-LeCENH3 (dilution 1:100) (Ma *et al*. 2016), rabbit anti-KNL1 (dilution 1:1000), rabbit anti-NDC80 (dilution 1:1000) (Oliveira *et al*. 2024) and mouse anti αtubulin (dilution 1:100, Sigma-Aldrich, St. Louis, MO; catalog number T6199). Antibodies against KNL1 and NDC80 were originally developed using respective peptides EDHFFGPVSPSFIRPGRLSDC and EQGINARDAERMKRELQALEG from *Cuscuta* sp. These epitopes have a respective 55.6% and 57.1% similarity to peptides DDNFFGPVSAKFLKSGRFSDT and EQEVNLRDVDRMKREMQLIER identified by tblastn similarity search of *Cuscuta europaea* KNL1 and NDC80 protein sequences in *L. sylvatica* genome. As the secondary antibody, goat anti-Rabbit IgG antibody (Invitrogen) or goat anti-mouse Alexa Fluor 488 (ImmunoResearch; catalog number 115-545-166) were used in a 1:500 dilution. Slides were incubated overnight at 4 °C, washed 3 times in 1×PBS and then the secondary antibody was applied, incubated at room temperature for 3h and washed 3 times in 1×PBS. The slides were counterstained with 2 µg/mL DAPI/Vectashield mounting medium. Microscopic images were recorded using a Zeiss Axiovert 200M microscope equipped with a Zeiss AxioCam CCD. Images were analyzed using the ZEN software (Carl Zeiss GmbH).

Oligo probes from the most abundant tandem repeats (*Lusy1* and *Lusy2*) and the *Arabidopsis* telomeric sequence (TTTAGGG) were used for fluorescent *in situ* hybridization (FISH). Mitotic chromosomes from roots pretreated with 2mM 8-hydroxyquinoline for 24 h at 4°C and fixed with ethanol:acetic acid (3:1 v/v) for 2 h, were prepared using the air-drying method (Ribeiro *et al*. 2016). FISH was performed as described by (Pedrosa *et al*. 2002). The slides were counterstained with 2 µg/mL DAPI in Vectashield (Vector) mounting medium. The images were captured as described above. Immuno-FISH was performed following (Houben *et al*. 2007), the immunostained slides were washed with 1xPBS for 15 min, postfixed in 4% paraformaldehyde in 1xPBS for 5 min, and then probed with the satellite *Lusy1*.

To analyze the chromatin ultrastructure, we applied super-resolution spatial structured illumination microscopy (3D-SIM) using a 63x/1.40 Oil Plan-Apochromat objective of an Elyra PS.1 microscope system and the ZENBlack software (Carl Zeiss GmbH) (Weisshart *et al*. 2016). Maximum intensity projections from image stacks were calculated from 3D-SIM image stacks. Zoom-in sections were presented as single slices to indicate the subnuclear chromatin structures at the super-resolution level.

## Supplementary Material

### Supplementary Figures

Figure S1: findGSE K-mer-based genome size estimation of *L. sylvatica*.

Figure S2: Detailed view of *L. sylvatica* chromosomes showing the dispersed distribution of the main genomic features.

Figure S3: *L. sylvatica* satellite characterization.

Figure S4: Proportion of CENH3 units (light blue), *Lusy1* (magenta) and *Lusy2* (green) arrays in 100kb windows.

Figure S5: Examples of (epi)genetic features distribution near satellite-based and satellite-free CENH3 domains of *L. sylvatica*.

Figure S6: Comparative analysis of GC content and epigenetic markers in functional/nonfunctional *Lusy* satellite arrays and satellite-free CENH3 domains.

Figure S7: Localization of CENH3, KNL1 and NDC80 in *Luzula nivea* (a) and *Juncus effusus* (b) metaphase chromosomes.

Figure S8: Phylogenetic relationships and comparative repeat analysis among *Luzula* species.

Figure S9: Genome synteny patterns showing macro-conserved blocks of *J. effusus* that are part of the *L. sylvatica* chromosomes.

Figure S10: *L. sylvatica* chromosomes showing the fusion of syntenic blocks of *J. effusus*.

Figure S11: Interstitial telomeric sites observed in the genome of *L. sylvatica*.

Figure S12: Characterization of fusion sites in *L. sylvatica* genome.

### Supplementary Tables

Table S1 Characteristics of the *L. sylvatica* genome assembly.

Table S2 Monomer sequence of the satellite DNA of *L. sylvatica* genome.

Table S3 Proportion of repeats in satellite-free CENH3 units of *L. sylvatica* genome.

Table S4 *Luzula* species, ENA codes and available genome size with their references used for comparative analyses.

Table S5 Individual genome proportion in percentage of the repetitive sequences in *Luzula* species.

Table S6 Comparative genome proportion of the repetitive sequences in *Luzula* species.

Table S7 Conserved syntenic blocks between *L. sylvatica* and *J. effusus* genome.

**Figure S1:**
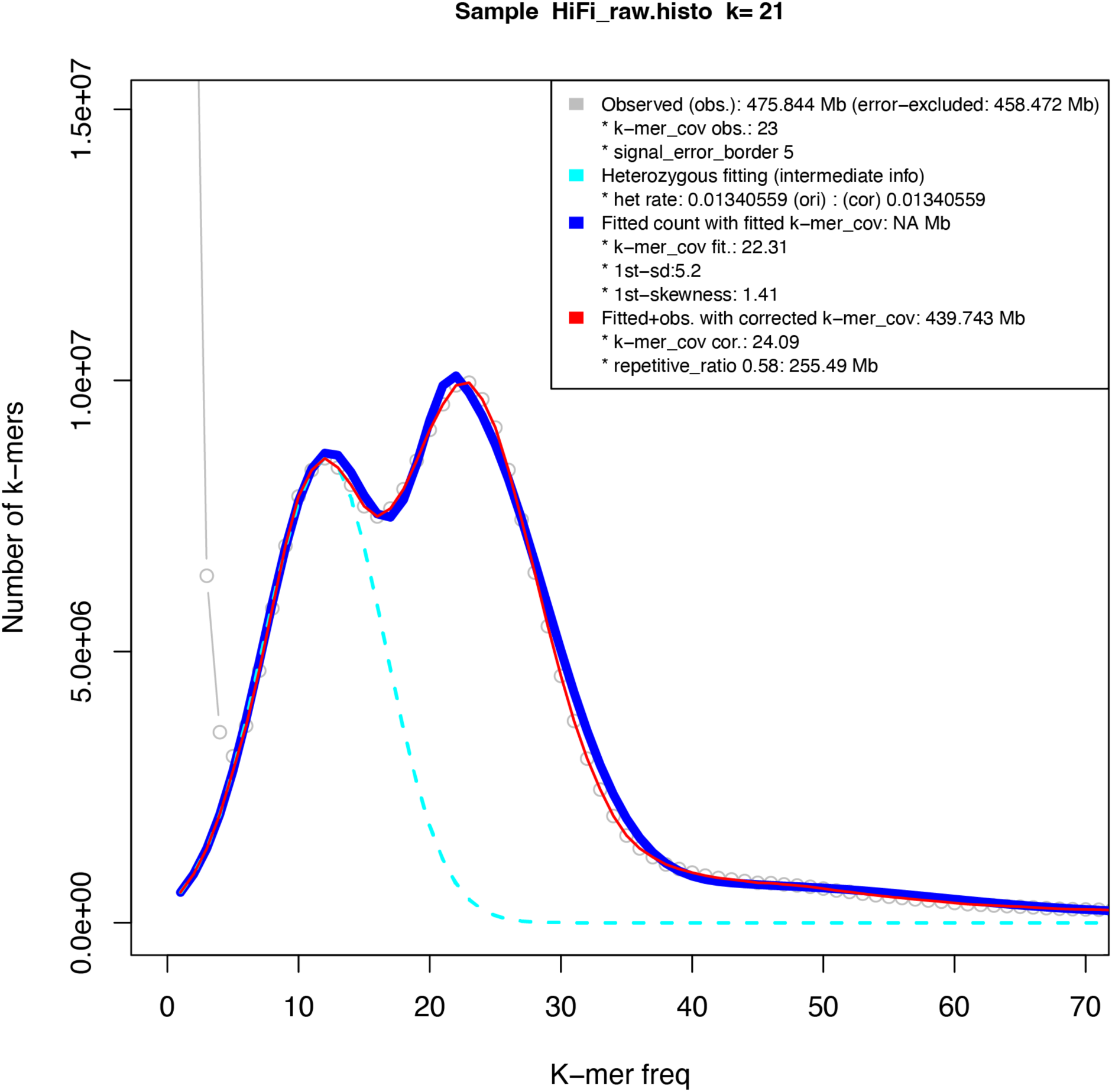
findGSE K-mer-based genome size estimation of *L. sylvatica*.

**Figure S2:**
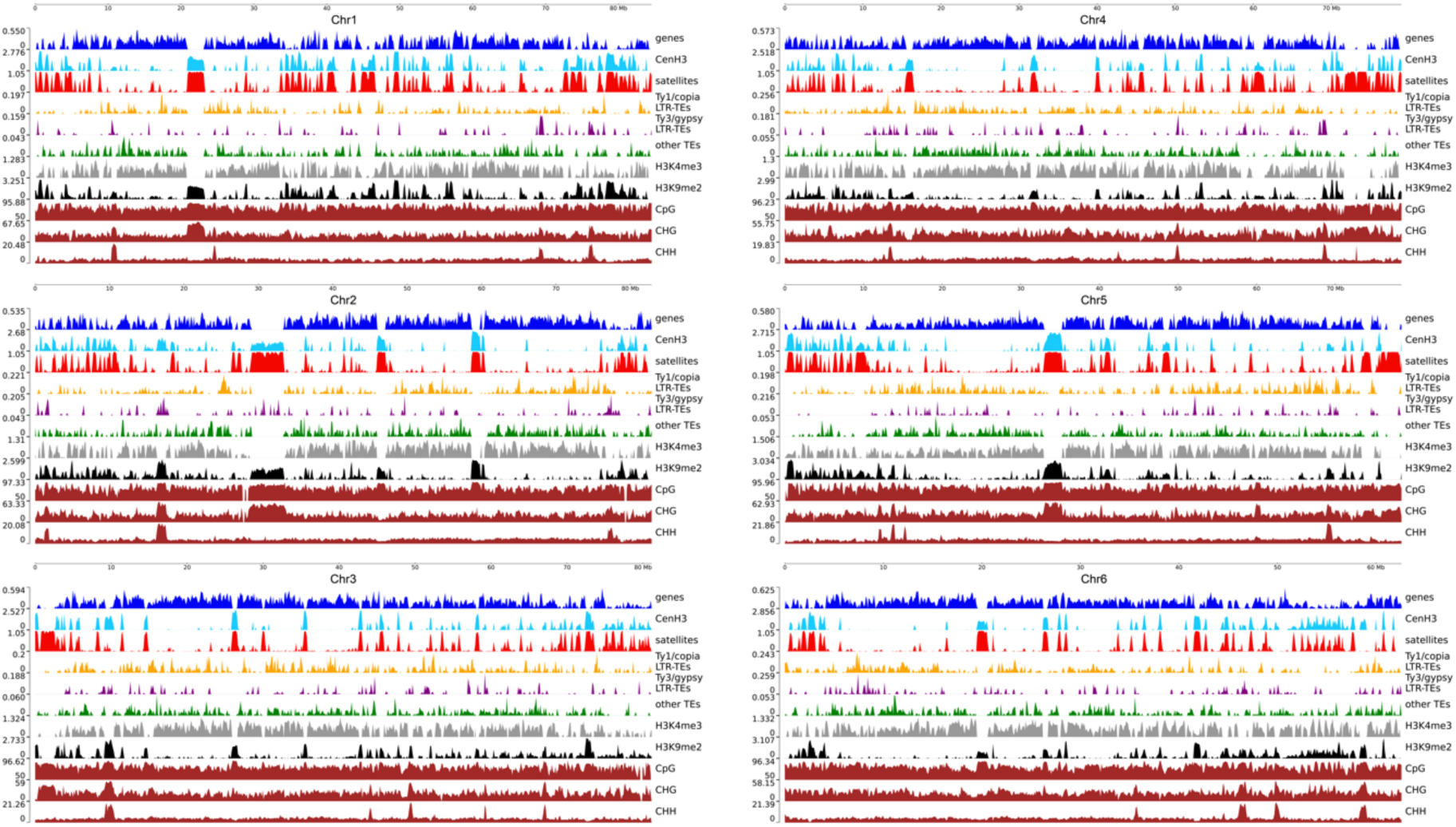
Detailed view of *L. sylvatica* chromosomes showing the dispersed distribution of the main genomic features. CENH3, gene, tandem repeat, dispersed repeat, eu-heterochromatin and histone mark densities, typical of holocentric chromosomes. Window sizes of 100 kb.

**Figure S3:**
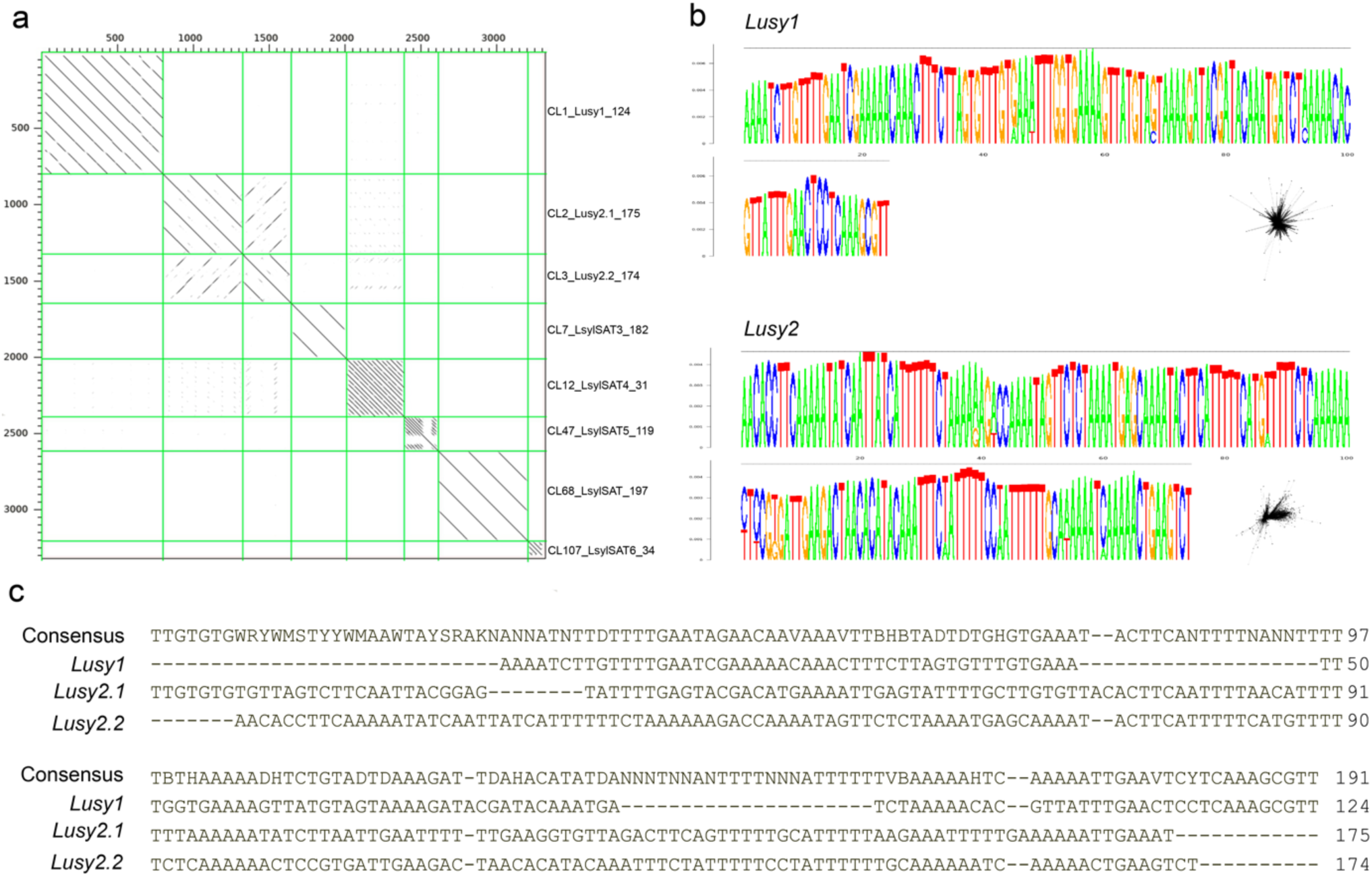
*L. sylvatica* satellite characterization. **(a)** Dot plot showing similarities between groups of tandem repeats. Despite being classified as a tandem pattern, LsylSAT_197 was not considered as a satellite because the genome mapping did not show a tandem distribution. **(b)** Sequence logo the most abundant satellite clusters LsylSatCL1 (*Lusy*1) and LsylSatCL2 (*Lusy*2). **(c)** Alignment between the two variants of *Lusy*2 (LsylvSat174 and LsylSat175) and *Lusy*1.

**Figure S4:**
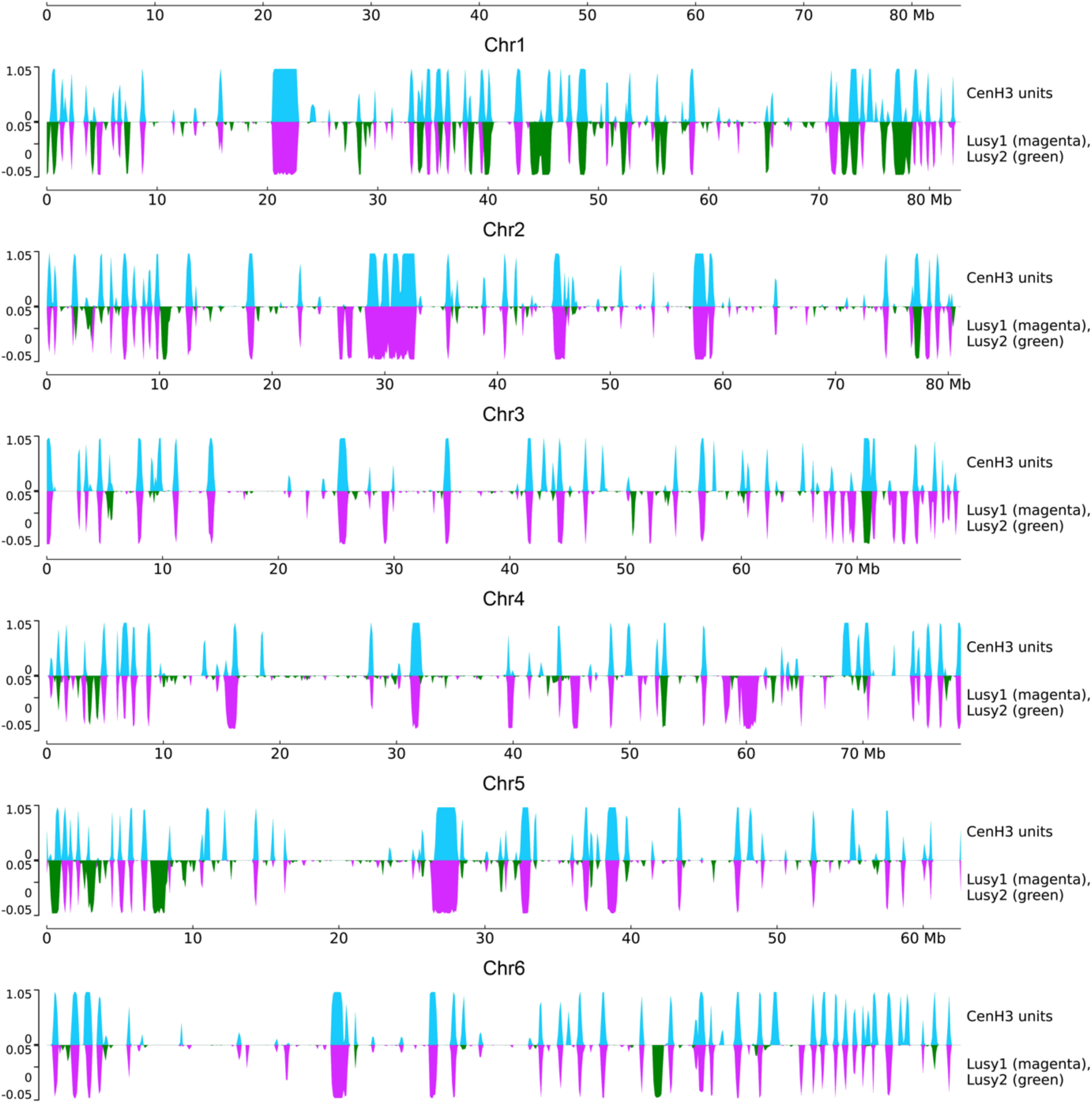
Proportion of CENH3 units (light blue), *Lusy1* (magenta) and *Lusy2* (green) arrays in 100kb windows.

**Figure S5:**
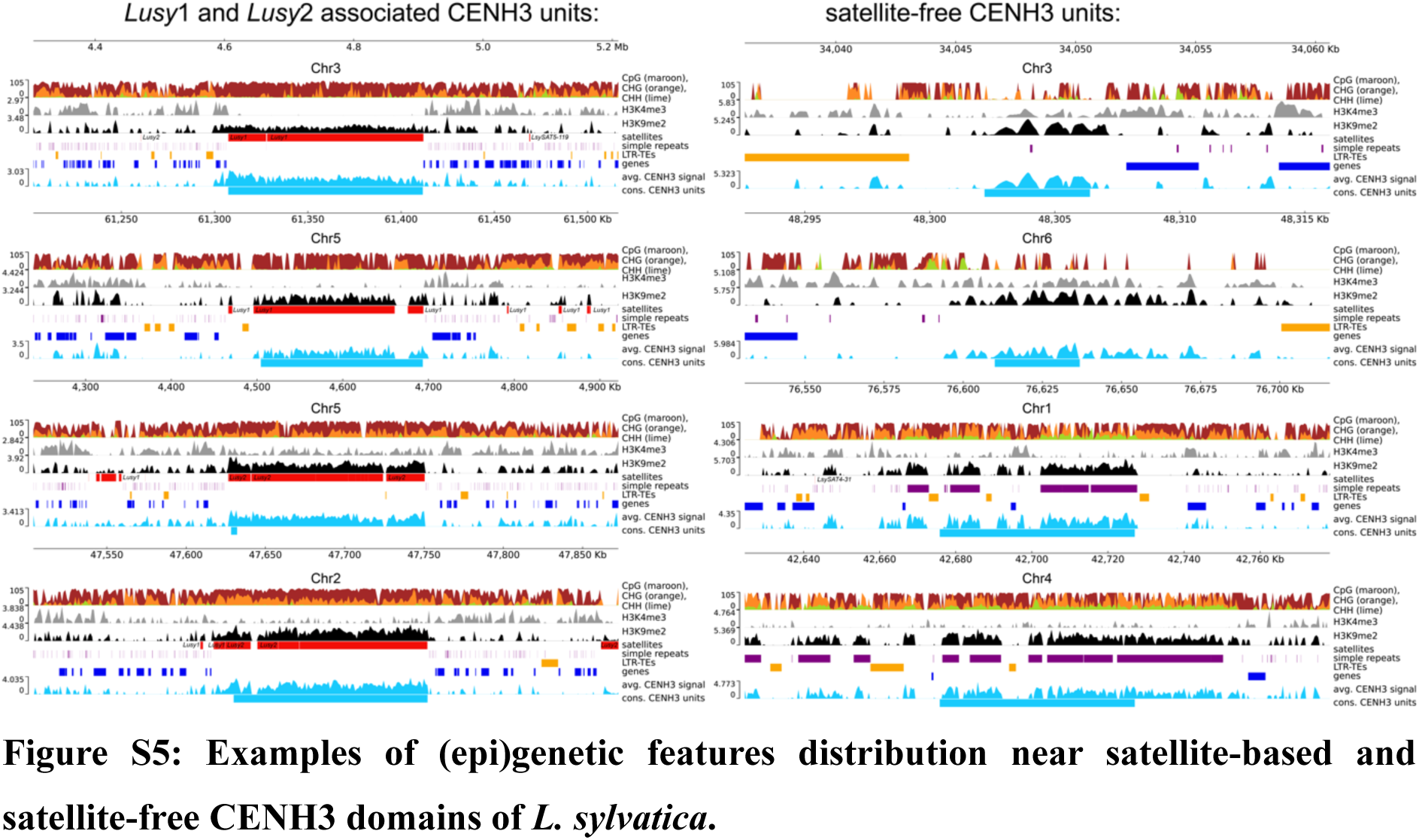
Examples of (epi)genetic features distribution near satellite-based and satellite-free CENH3 domains of *L. sylvatica*.

**Figure S6:**
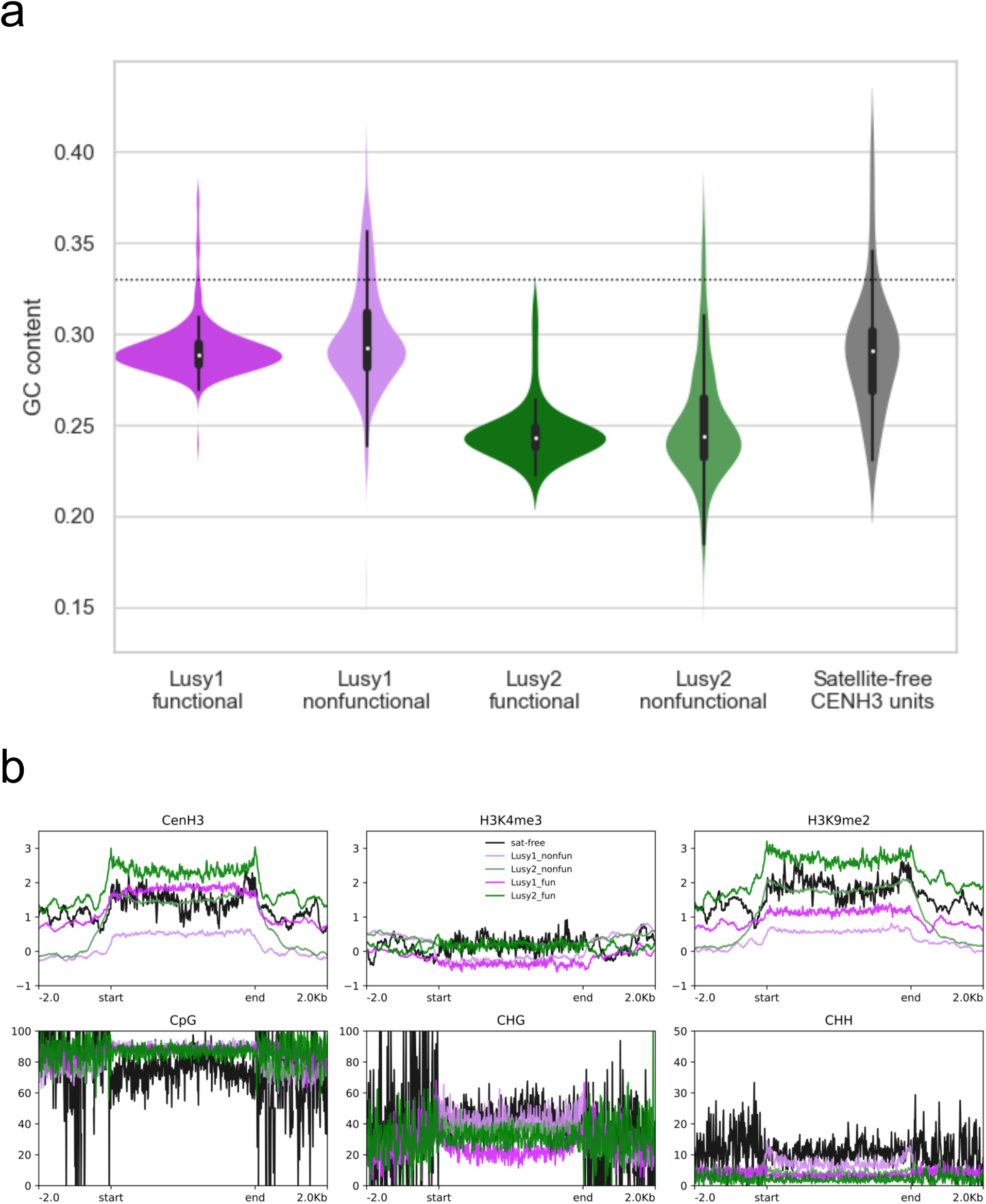
Comparative analysis of GC content and epigenetic markers in functional/nonfunctional *Lusy* satellite arrays and satellite-free CENH3 domains. (**a**) GC content of functional and nonfunctional satellite arrays and satellite-free CENH3 units. Dotted line at the overall genomic GC content level. (**b**) Metaplots showing the level of enrichment of functional and nonfunctional satellite arrays and *Lusy* satellite-free CENH3 units with CENH3, H3K4me3, H3K9me2, CpG, CHH, and CGH.

**Figure S7:**
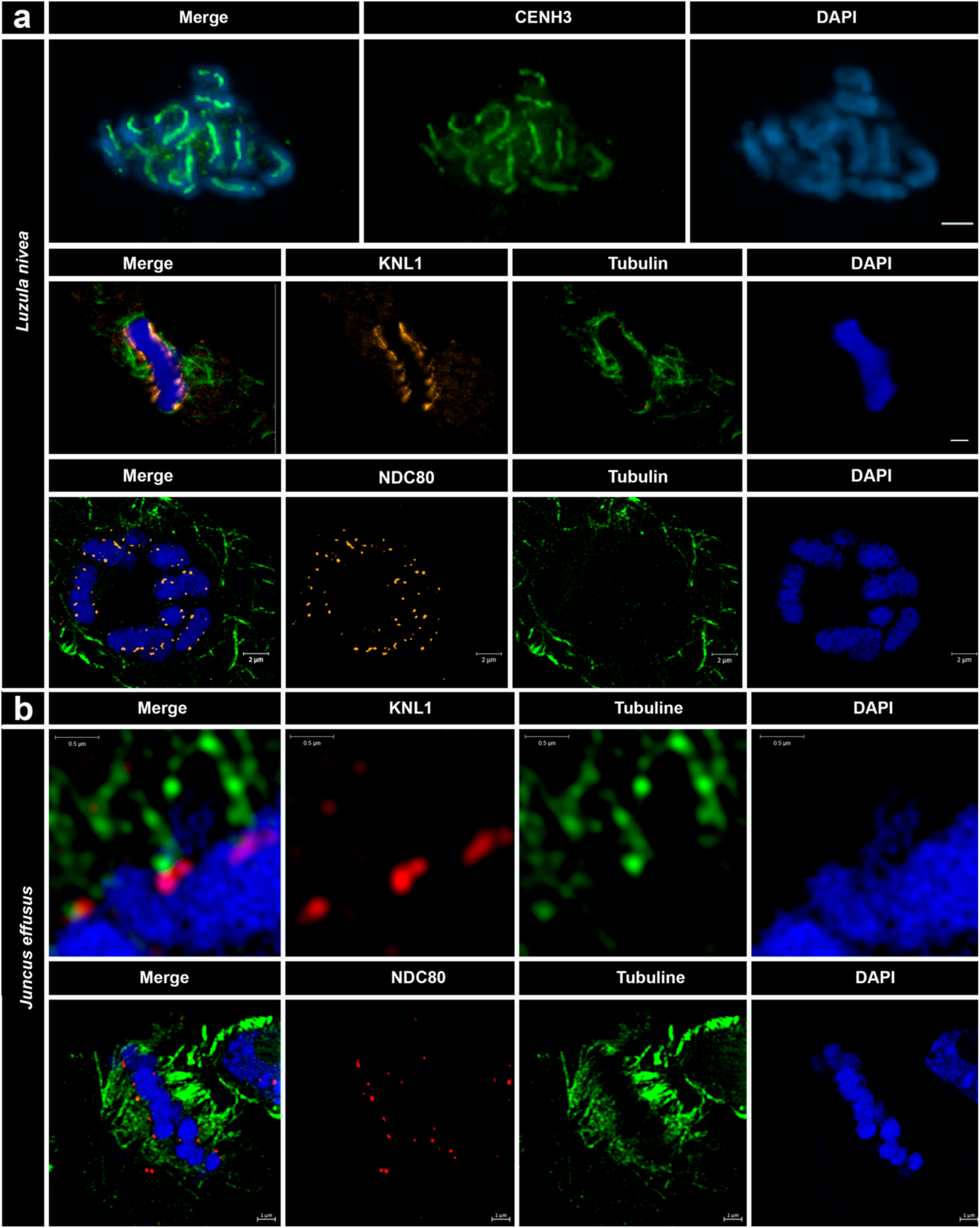
Localization of CENH3, KNL1 and NDC80 in *Luzula nivea* (a) and *Juncus effusus* (b) metaphase chromosomes. KNL1 and NDC80 localize specifically to the surface of the *J. effusus* centromere, where microtubules attach. The images in **(b)** represent super-resolution (a single slice of a 3D-SIM image stack.

**Figure S8:**
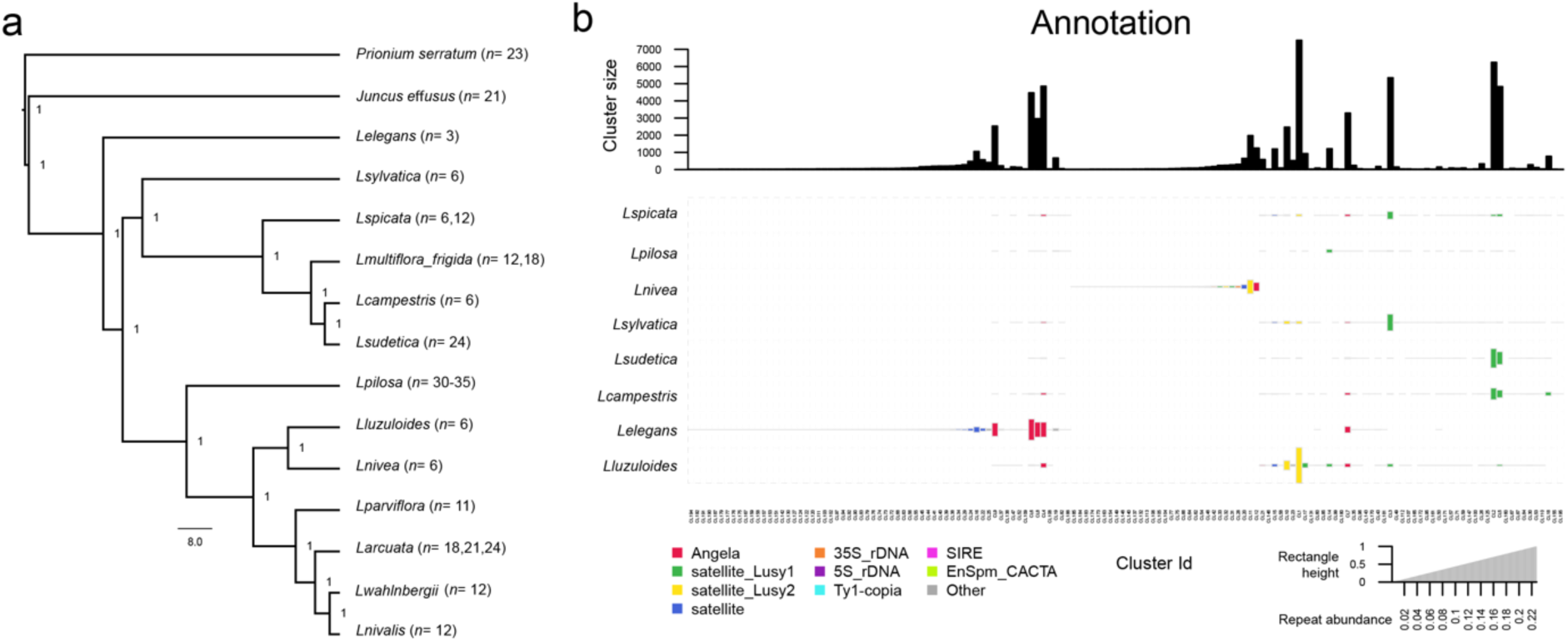
Phylogenetic relationships and comparative repeat analysis among *Luzula* species. (**a**) Whole-plastome phylogenetic analysis. The plastome of *Juncus effusus* (MW366789) and *Prionium serratum* (OL689155) were used as outgroup. Chromosome numbers were recorded here and compiled from Záveská Drábková (2013). (**b**) Comparative abundance of the main types of repetitive sequences in *Luzula* species. The size of the rectangle is proportional to the genome abundance of that cluster for each species. The colors of the balls correspond to different repetitive sequence types. The proportion of each cluster was adjusted according to genome size.

**Figure S9:**
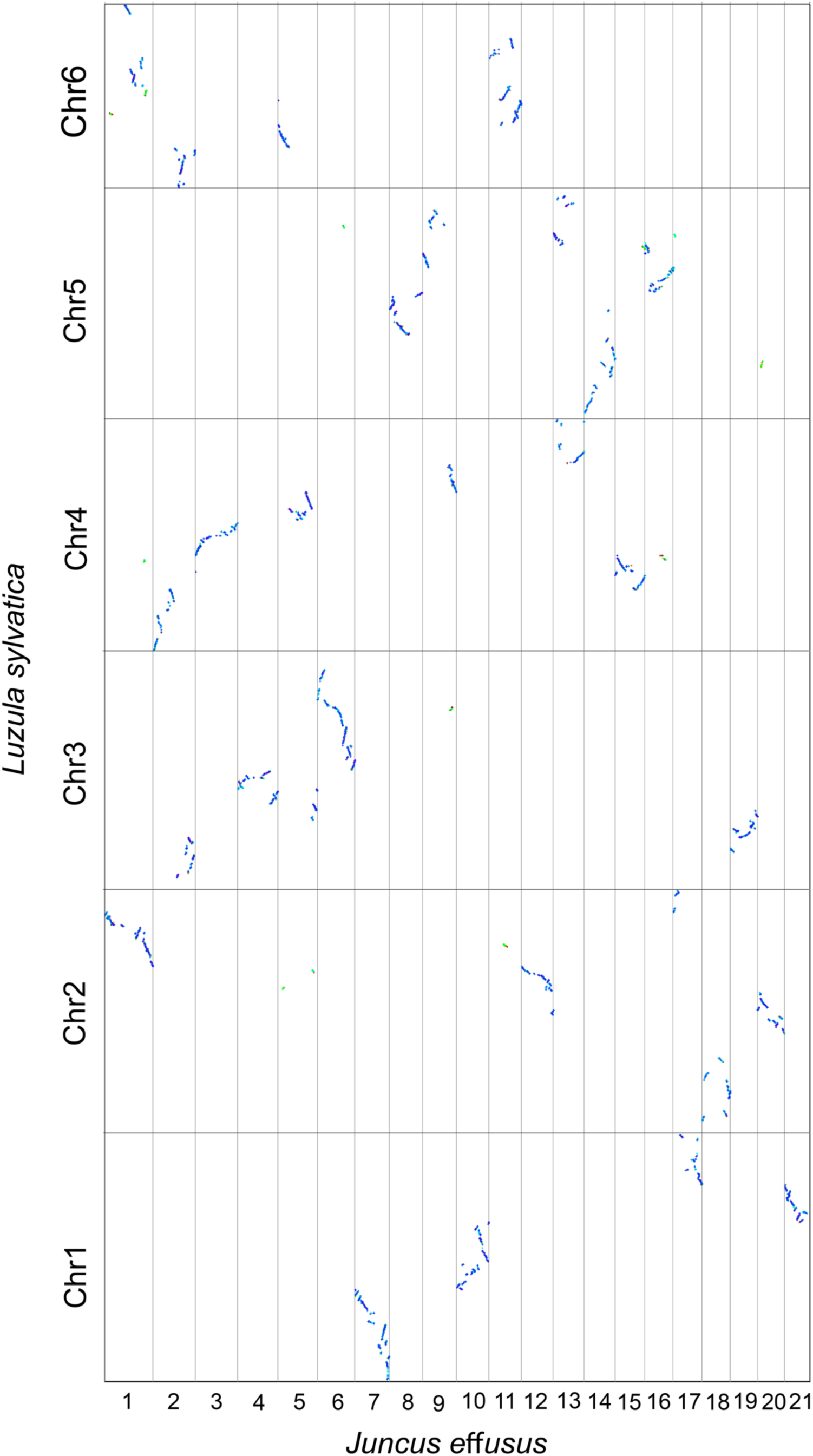
Genome synteny patterns showing macro-conserved blocks of *J. effusus* that are part of the *L. sylvatica* chromosomes.

**Figure S10:**
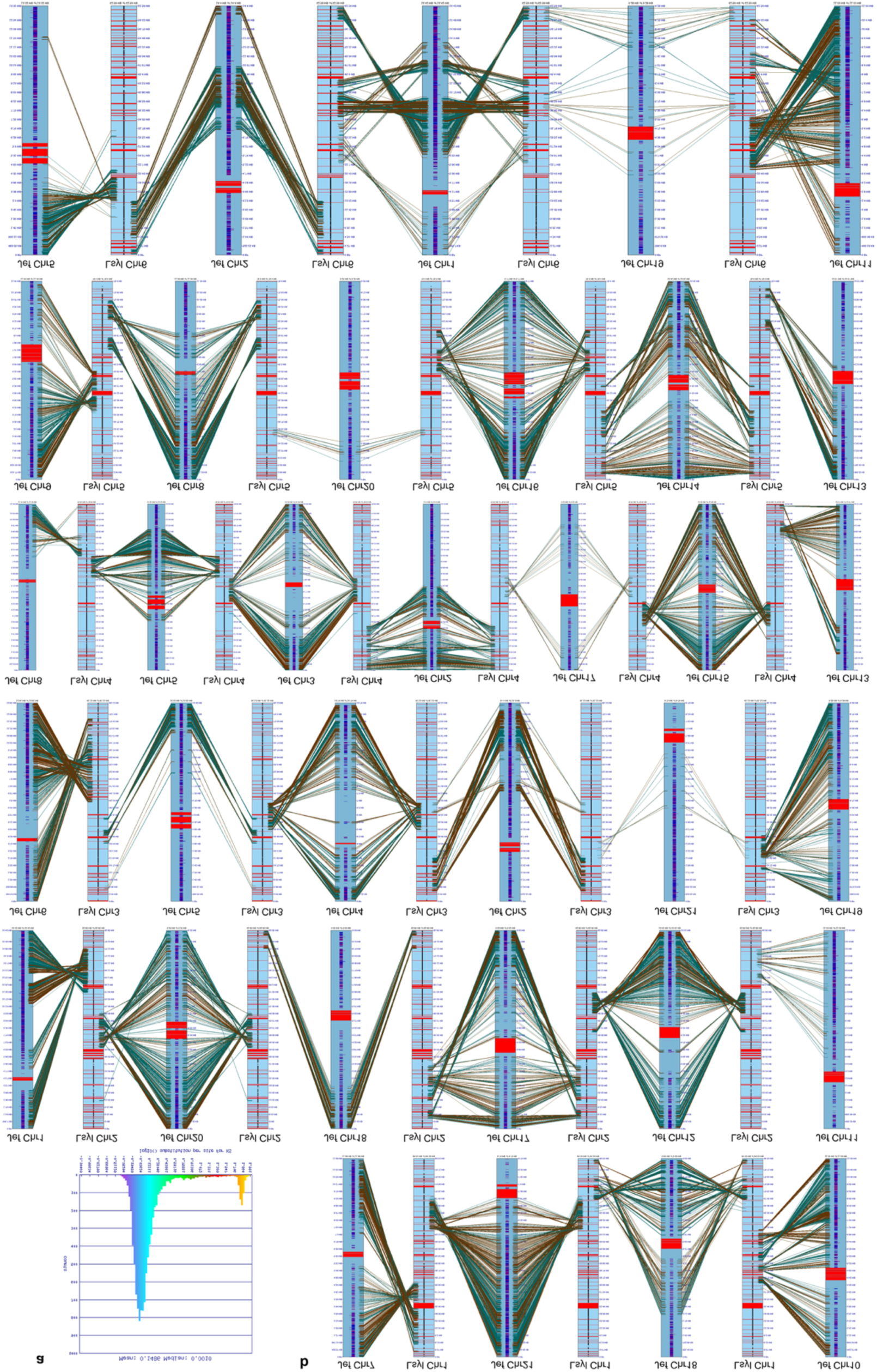
*L. sylvatica* chromosomes showing the fusion of syntenic blocks of *J. effusus*. **(a)** Synonymous substitution rate (Ks) determined for *L. sylvatica* genome using CodeML. **(b)** The synteny stops near the centromeric CENH3 (green) sites. Genes, CENH3 and telomere domains are annotated as blue-purple, green and black stripes, respectively.

**Figure S11:**
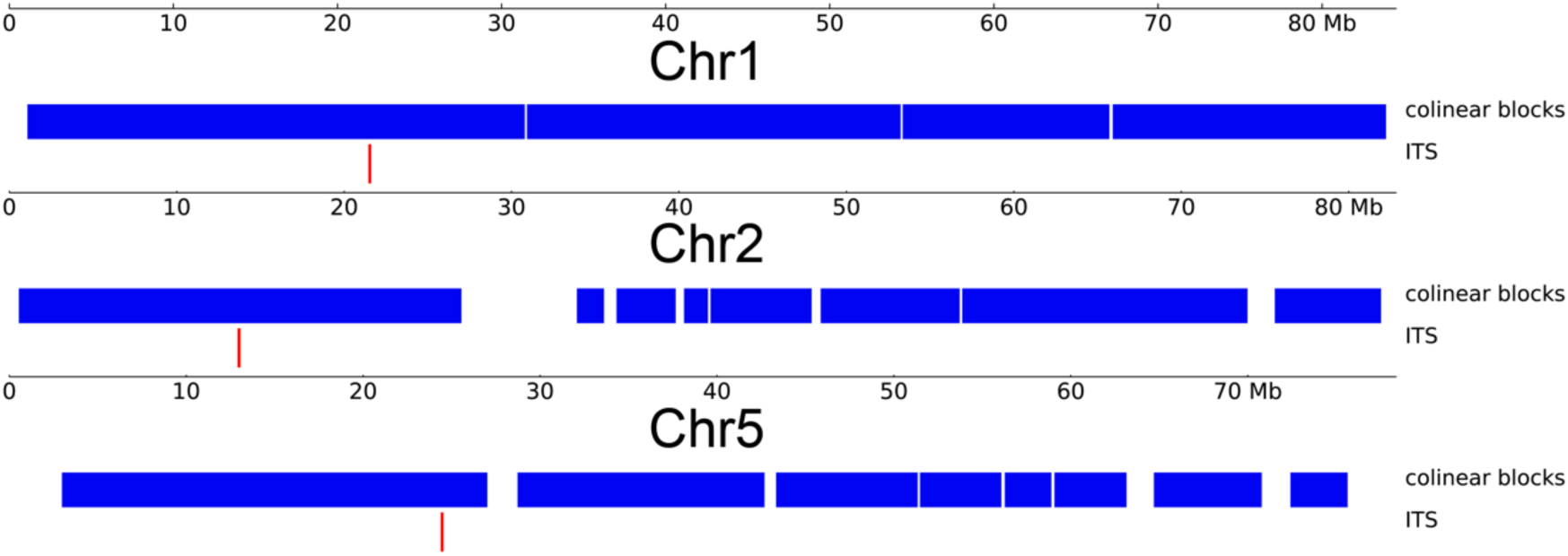
Interstitial telomeric sites observed in the genome of *L. sylvatica*. Blue blocks show the syntenic regions with the *Juncus effusus* genome. Interstitial sites are nowhere near the fusion points.

**Figure S12:**
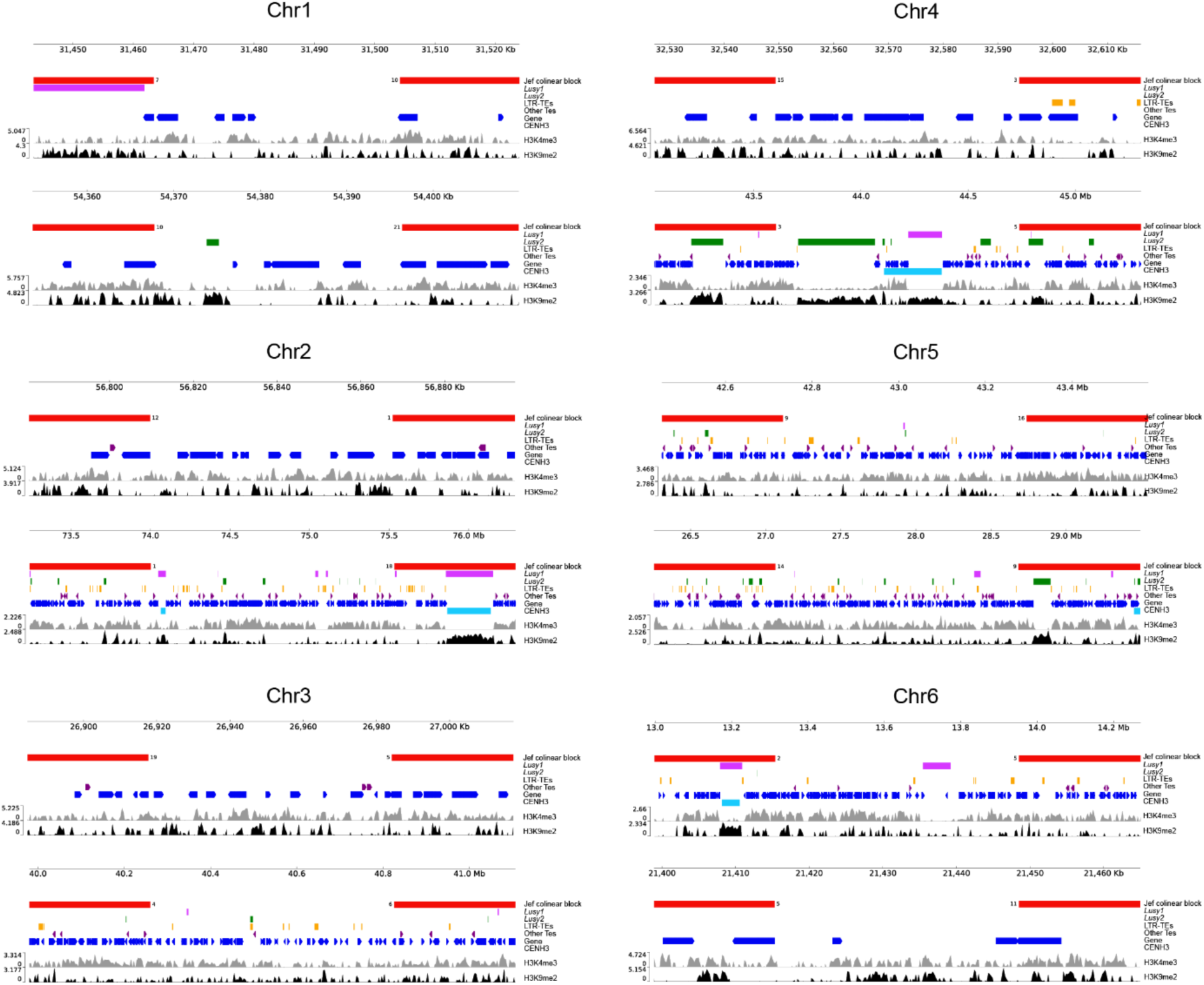
Characterization of fusion sites in *L. sylvatica* genome. Similar fusion signatures are shared between some chromosomes, where regions enriched with gene, TE or *Lusy2* repeats are located either up or downstream of the fusion point.

**Table S1.**
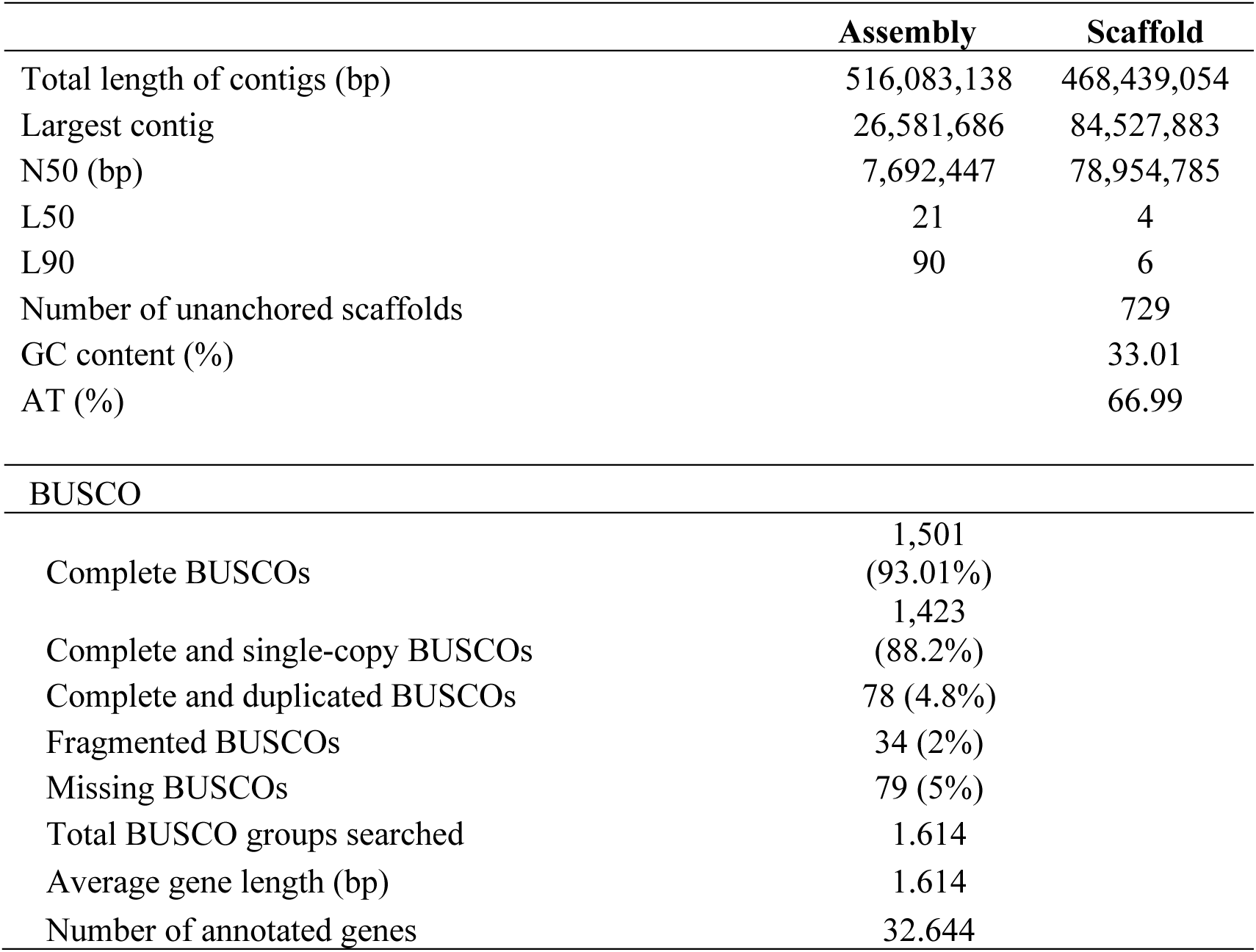
Characteristics of the *Luzula sylvatica* genome assembly.

**Table S2.**
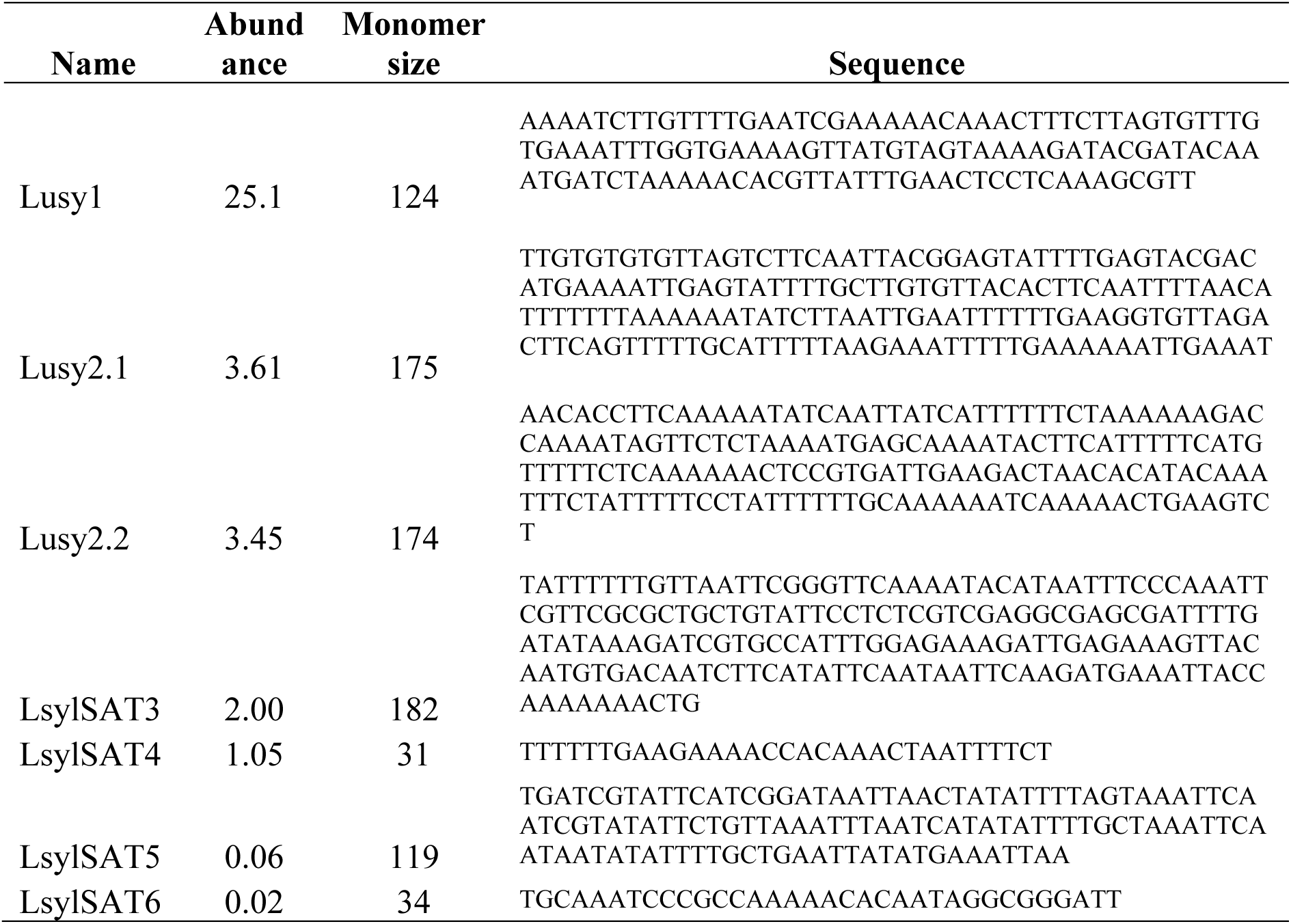
Monomer sequence of the satellite DNA of *Luzula sylvatica* genome.

**Table S3.** Proportion of repeats in satellite-free CENH3 units of *L. sylvatica* genome.

| **Name** | **count in**<br>**sat-free**<br>**CENH3**<br>**domains** | **Total length in**<br>**sat-free CENH3**<br>**domains** | **proportion**<br>**of sat-free**<br>**CENH3**<br>**domains**<br>**(%)** | **proportion**<br>**in genome**<br>**(%)** |
| --- | --- | --- | --- | --- |
| Class_I/LTR/Ty3-gypsy/non-chromovirus/OTA/Athila | 392 | 582,139.00 | 18 | 2.6 |
| Class_I/LTR/Ty1-copia/Angela | 215 | 158,332.00 | 5 | 10 |
| Class_I/LTR/Ty1-copia/SIRE | 58 | 49,920.00 | 2 | 1.6 |
| Class_I/LTR/Ty1-copia/Bianca | 48 | 21,209.00 | 1 | 0.7 |
| Class_I/LTR/Ty3-gypsy/chromovirus/Tekay | 15 | 13,567.00 | 0 | 0.2 |
| Class_I/LTR/Ty1-copia/Tork | 11 | 10,678.00 | 0 | 0.3 |
| Class_I/LTR/Ty3-gypsy/chromovirus/Reina | 7 | 5,811.00 | 0 | 0.2 |
| Class_I/LTR/Ty1-copia/Ale | 9 | 4,577.00 | 0 | 0.4 |
| Class_I/LTR/Ty1-copia/Ivana | 13 | 2,676.00 | 0 | 1.0 |
| Class_I/LTR/Ty1-copia/TAR | 2 | 46.00 | 0 | 0.1 |
| Low_complexity | 88 | 4,628.00 | 0 | 0.4 |
| Simple_repeat | 643 | 1,782,330.00 | 54 | 3.7 |
| Unknown | 2 | 4.00 | 0 | 0 |

**Table S4.**
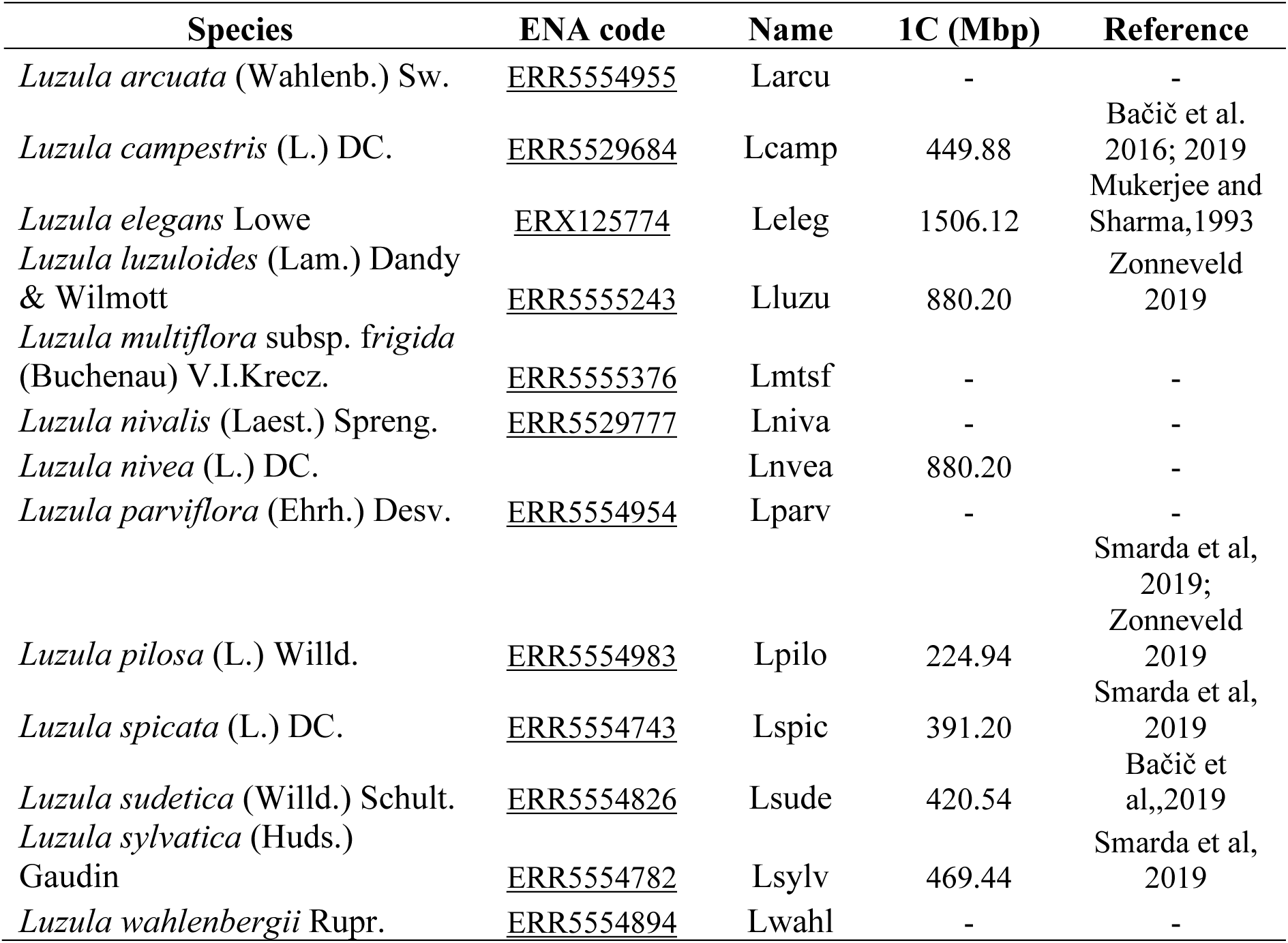
*Luzula* species, ENA codes and available genome size with their references used for comparative analyses.

**Table S5.** Individual genome proportion in percentage of the repetitive sequences in *Luzula* species.

|  | Larcu | Lwahl | Lluzu | Lparv | Lniva | Lsylv | Lspic | Lcamp | Lsude | Lmtsf | Lpilo | Lnvea | Leleg |
| --- | --- | --- | --- | --- | --- | --- | --- | --- | --- | --- | --- | --- | --- |
| **Tandem** |  |  |  |  |  |  |  |  |  |  |  |  |  |
| 35S_rDNA | 0.72 | 0.74 | 0.38 | 0.93 | 0.89 | 1.34 | 0.53 | 0.79 | 0.53 | 0.85 | 1.02 | 2.14 | 0.04 |
| 5S_rDNA | 0.08 | 0.21 | 0.06 | 0.12 | 0.07 | 0.09 | 0.09 | 0.02 | - | 0.02 | 0.06 | 0.16 | 0.21 |
| Satellite_Lusy1 | 15.89 | 22.10 | 9.44 | 12.82 | 13.66 | 25.1 | 20.02 | 33.68 | 47.41 | 23.65 | 13.35 | 3.33 | - |
| Satellite_Lusy2 | 10.12 | 20.28 | 31.03 | 7.95 | 4.44 | 7.08 | 5.41 | 0.43 | 0.76 | 0.34 | - | 15.54 | 0.72 |
| Other Satellites | 1.69 | 1.67 | 4.07 | 1.87 | 1.45 | 3.15 | 2.56 | 0.78 | 0.78 | 1.18 | 18.72 | 1.21 | 9.69 |
| **LTR** |  |  |  |  |  |  |  |  |  |  |  |  |  |
| Ty1_copia/Alesia | 0.13 | 0.05 | - | 0.11 | 0.16 | - | - | - | - | - | - | 0.03 | 0.02 |
| Ty1_copia/Angela | 21.49 | 16.04 | 8.84 | 25.45 | 22.5 | 5.87 | 10.49 | 10.85 | 3.69 | 12.49 | 1.21 | 12.36 | 37.61 |
| Ty1_copia/Bianca | - | - | - | 0.2 | 0.31 | 0.03 | 0.02 | 0.15 | 0.03 | 0.27 | - | 0.12 | - |
| Ty1_copia/Ivana | 0.26 | 0.14 | 0.02 | 0.34 | 0.38 | 0.13 | 0.06 | 0.02 | - | 0.04 | - | 0.19 | 2.26 |
| Ty1_copia/SIRE | 3.25 | 2.02 | 0.07 | 4.18 | 4.52 | 1.37 | 2.1 | 0.15 | - | 1.64 | - | 2.14 | 1.46 |
| Ty1_copia/Tork | 0.03 | - | - | 0.07 | 0.17 | 0.13 | - | - | - | - | - | 0.2 | - |
| Ty3_gypsy/chromovirus/Tekay | - | 0.02 | 0.05 | 0.38 | 0.29 | - | - | - | - | - | - | 1.1 | 0.94 |
| Ty3_gypsy/non-chromovirus/Athila | 1.43 | - | - | 0.16 | 1.78 | 0.27 | 0.07 | 0.11 | - | 0.17 | - | 0.04 | - |
| **No-LTR** |  |  |  |  |  |  |  |  |  |  |  |  |  |
| LINE | - | - | - | - | - | - | - | - | - | - | - | - | 0.3 |
| Pararetrovirus | - | - | - | - | - | - | - | - | - | - | - | 0.01 | - |
| TIR/EnSpm_CACTA | 2.34 | 0.03 | 0.03 | 1.72 | 2.02 | - | - | - | - | - | - | 2.19 | 2.16 |
| TIR/hAT | 0.29 | - | - | - | - | - | - | - | - | - | - | - | 0.03 |
| TIR/MuDR_Mutator | - | - | - | 0.04 | 0.03 | - | - | - | - | - | - | - | 0.22 |
| Unclassified | 2.78 | 2.99 | 4.89 | 4.44 | 3.34 | 3.67 | 2.68 | 2.43 | 1.71 | 3.72 | 0.88 | 1.91 | 7.79 |
| **Total** | 60.49 | 66.29 | 58.88 | 60.78 | 56.01 | 48.23 | 44.03 | 49.41 | 54.91 | 44.37 | 35.24 | 42.67 | 63.45 |

**Table S6.** Comparative genome proportion of the repetitive sequences in *Luzula* species.

| Elem | cluster | Larcu | Lwahl | Lluzu | Lparv | Lniva | Lsylv | Lspic | Lcamp | Lsude | Lmtsf | Lpilo | Lnvea | Leleg |
| --- | --- | --- | --- | --- | --- | --- | --- | --- | --- | --- | --- | --- | --- | --- |
| Lusy2 | 1 | 14.85 | 14.59 | 24.20 | 5.56 | 3.23 | 5.78 | 3.88 | 0.45 | 0.63 | 0.28 | 0.01 | 1.22 | 0.00 |
| Lusy1 | 2 | 6.62 | 8.68 | 2.00 | 4.00 | 3.53 | 17.10 | 10.31 | 0.32 | 0.23 | 0.48 | 1.71 | 0.00 | 0.00 |
| Ty1-copia/Angela | 3 | 11.74 | 4.07 | 2.75 | 7.08 | 6.34 | 1.57 | 2.94 | 2.45 | 0.89 | 2.90 | 0.48 | 2.78 | 5.06 |
| Ty1-copia/Angela | 4 | 10.88 | 3.80 | 2.21 | 6.51 | 5.88 | 1.59 | 2.68 | 2.43 | 1.08 | 3.06 | 0.49 | 2.30 | 4.29 |
| Lusy1 | 5 | 2.98 | 3.18 | 0.74 | 1.82 | 1.31 | 0.42 | 3.01 | 8.04 | 14.03 | 5.22 | 0.67 | 0.00 | 0.00 |
| Lusy1 | 6 | 0.02 | 0.00 | 0.02 | 0.02 | 0.00 | 0.02 | 1.38 | 8.57 | 14.84 | 5.45 | 0.02 | 0.00 | 0.00 |
| Ty1-copia/Angela | 7 | 6.74 | 2.20 | 1.71 | 3.72 | 3.55 | 0.96 | 1.60 | 1.42 | 0.70 | 1.96 | 0.23 | 0.00 | 5.57 |
| Lusy1 | 8 | 7.51 | 3.06 | 1.79 | 2.55 | 3.27 | 0.00 | 0.00 | 0.00 | 0.00 | 0.00 | 8.55 | 0.00 | 0.00 |
| Lusy1 | 9 | 0.04 | 0.08 | 0.04 | 0.02 | 0.02 | 0.01 | 0.99 | 5.30 | 8.31 | 3.15 | 0.06 | 0.00 | 0.00 |
| Lusy1 | 10 | 6.93 | 2.08 | 2.97 | 1.82 | 2.81 | 0.00 | 0.00 | 0.01 | 0.00 | 0.01 | 0.08 | 0.00 | 0.00 |
| All/repeat/satellite | 11 | 1.34 | 0.82 | 1.40 | 0.62 | 0.91 | 1.76 | 1.80 | 0.61 | 0.48 | 0.76 | 0.01 | 0.00 | 0.00 |
| Ty1-copia/Angela | 12 | 1.64 | 0.88 | 0.75 | 1.83 | 1.21 | 0.42 | 0.76 | 0.65 | 0.37 | 1.03 | 0.22 | 0.00 | 0.00 |
| Lusy1 | 14 | 0.01 | 0.00 | 0.00 | 0.00 | 0.00 | 0.00 | 0.19 | 4.19 | 0.27 | 3.12 | 0.00 | 0.07 | 0.00 |
| 35S_rDNA | 19 | 0.62 | 0.33 | 0.15 | 0.31 | 0.24 | 0.47 | 0.15 | 0.17 | 0.18 | 0.22 | 0.17 | 0.09 | 0.02 |
| All/repeat/satellite | 23 | 0.30 | 0.19 | 0.08 | 0.20 | 0.21 | 0.18 | 0.14 | 0.08 | 0.15 | 0.10 | 0.40 | 0.00 | 0.00 |
| All/repeat/satellite | 24 | 0.00 | 0.00 | 0.00 | 0.00 | 0.00 | 0.00 | 0.00 | 0.00 | 0.00 | 0.00 | 0.00 | 0.00 | 2.02 |
| Ty1-copia/Angela | 25 | 0.00 | 0.00 | 0.00 | 0.00 | 0.00 | 0.00 | 0.00 | 0.00 | 0.00 | 0.00 | 0.00 | 0.00 | 1.74 |
| All/repeat/satellite | 26 | 0.04 | 0.02 | 0.04 | 0.12 | 0.11 | 0.13 | 0.24 | 0.04 | 0.06 | 0.13 | 0.46 | 0.00 | 0.22 |
| 35S_rDNA | 28 | 0.14 | 0.09 | 0.05 | 0.09 | 0.06 | 0.24 | 0.09 | 0.14 | 0.10 | 0.16 | 0.22 | 0.00 | 0.01 |
| All/repeat/satellite | 34 | 0.00 | 0.00 | 0.00 | 0.00 | 0.00 | 0.00 | 0.00 | 0.00 | 0.00 | 0.00 | 0.00 | 0.00 | 0.94 |
| All/repeat/satellite | 35 | 0.00 | 0.00 | 0.00 | 0.00 | 0.00 | 0.85 | 0.00 | 0.00 | 0.00 | 0.00 | 0.00 | 0.00 | 0.00 |
| All/repeat/satellite | 36 | 0.00 | 0.00 | 0.00 | 0.00 | 0.00 | 0.00 | 0.00 | 0.00 | 0.00 | 0.00 | 0.00 | 0.00 | 0.86 |
| 35S_rDNA | 37 | 0.24 | 0.11 | 0.06 | 0.06 | 0.06 | 0.09 | 0.03 | 0.02 | 0.04 | 0.09 | 0.00 | 0.00 | 0.02 |
| All/repeat/satellite | 38 | 0.00 | 0.00 | 0.00 | 0.00 | 0.00 | 0.00 | 0.00 | 0.00 | 0.00 | 0.00 | 0.00 | 0.00 | 0.81 |
| 5S_rDNA | 41 | 0.13 | 0.18 | 0.03 | 0.11 | 0.05 | 0.05 | 0.03 | 0.02 | 0.00 | 0.00 | 0.01 | 0.00 | 0.10 |
| Ty1-copia/SIRE | 46 | 0.20 | 0.09 | 0.02 | 0.09 | 0.06 | 0.02 | 0.02 | 0.02 | 0.00 | 0.03 | 0.00 | 0.11 | 0.06 |
| Ty1-copia/Angela | 49 | 0.00 | 0.00 | 0.00 | 0.00 | 0.00 | 0.00 | 0.00 | 0.00 | 0.00 | 0.00 | 0.00 | 0.18 | 0.58 |
| Lusy1 | 60 | 0.04 | 0.11 | 0.03 | 0.03 | 0.00 | 0.02 | 0.01 | 0.07 | 0.01 | 0.07 | 0.00 | 0.13 | 0.00 |
| Ty1-copia | 61 | 1.25 | 0.54 | 0.39 | 0.96 | 0.89 | 0.12 | 0.56 | 0.83 | 0.17 | 0.67 | 0.05 | 0.00 | 0.00 |
| Ty1-copia/SIRE | 69 | 0.11 | 0.02 | 0.02 | 0.07 | 0.09 | 0.02 | 0.02 | 0.01 | 0.01 | 0.01 | 0.00 | 0.00 | 0.02 |
| All/repeat/satellite | 72 | 0.00 | 0.00 | 0.00 | 0.00 | 0.00 | 0.00 | 0.00 | 0.00 | 0.00 | 0.00 | 0.00 | 0.00 | 0.37 |
| All/repeat/satellite | 78 | 0.00 | 0.00 | 0.00 | 0.00 | 0.00 | 0.03 | 0.00 | 0.00 | 0.00 | 0.00 | 0.02 | 0.00 | 0.27 |
| Ty1-copia/Angela | 79 | 0.00 | 0.00 | 0.00 | 0.00 | 0.00 | 0.00 | 0.00 | 0.00 | 0.00 | 0.00 | 0.00 | 0.00 | 0.33 |
| Ty1-copia/Angela | 89 | 0.00 | 0.00 | 0.00 | 0.00 | 0.00 | 0.00 | 0.00 | 0.00 | 0.00 | 0.00 | 0.00 | 0.00 | 0.28 |
| Ty1-copia/SIRE | 93 | 0.07 | 0.02 | 0.03 | 0.05 | 0.05 | 0.02 | 0.01 | 0.00 | 0.00 | 0.01 | 0.00 | 0.00 | 0.00 |
| Ty1-copia/Angela | 104 | 0.03 | 0.05 | 0.01 | 0.10 | 0.01 | 0.00 | 0.00 | 0.01 | 0.01 | 0.01 | 0.00 | 0.08 | 0.00 |
| Ty1-copia/Ivana | 106 | 0.09 | 0.02 | 0.00 | 0.03 | 0.02 | 0.01 | 0.03 | 0.00 | 0.01 | 0.02 | 0.00 | 0.00 | 0.00 |
| Lusy2 | 109 | 0.00 | 0.00 | 0.00 | 0.00 | 0.00 | 0.00 | 0.00 | 0.00 | 0.00 | 0.00 | 0.00 | 0.00 | 0.24 |
| All/repeat/satellite | 112 | 0.00 | 0.05 | 0.00 | 0.16 | 0.02 | 0.00 | 0.00 | 0.00 | 0.00 | 0.00 | 0.00 | 0.00 | 0.00 |
| Ty1-copia/SIRE | 113 | 0.07 | 0.02 | 0.05 | 0.03 | 0.03 | 0.00 | 0.02 | 0.00 | 0.00 | 0.01 | 0.00 | 0.00 | 0.00 |
| Ty1-copia/Angela | 122 | 0.02 | 0.01 | 0.01 | 0.02 | 0.02 | 0.02 | 0.05 | 0.01 | 0.03 | 0.02 | 0.00 | 0.00 | 0.01 |
| Ty1-copia/SIRE | 128 | 0.07 | 0.00 | 0.02 | 0.02 | 0.03 | 0.01 | 0.00 | 0.03 | 0.00 | 0.01 | 0.01 | 0.00 | 0.00 |
| Ty1-copia/Ivana | 130 | 0.04 | 0.00 | 0.03 | 0.03 | 0.01 | 0.02 | 0.00 | 0.01 | 0.01 | 0.04 | 0.01 | 0.00 | 0.00 |
| All/repeat/satellite | 138 | 0.00 | 0.00 | 0.00 | 0.19 | 0.00 | 0.00 | 0.00 | 0.00 | 0.00 | 0.00 | 0.00 | 0.00 | 0.00 |
| All/repeat/satellite | 161 | 0.16 | 0.00 | 0.00 | 0.00 | 0.00 | 0.00 | 0.00 | 0.00 | 0.00 | 0.00 | 0.00 | 0.00 | 0.00 |

**Table S7.** Conserved collinear blocks between *Luzula sylvatica* and *Juncus effusus* genome.

| *Juncus effusus*<br>Chr | Start | End | *Luzula sylvatica*<br>Chr | Start | End |
| --- | --- | --- | --- | --- | --- |
| 1 | 51433 | 3183432 | 3 | 70363021 | 75329258 |
| 1 | 10100862 | 16389583 | 3 | 56270627 | 70062240 |
| 1 | 5428881 | 6590660 | 3 | 69713915 | 70969951 |
| 1 | 4955851 | 5219128 | 3 | 70246702 | 70357775 |
| 1 | 6419584 | 8660074 | 4 | 59222634 | 62351670 |
| 1 | 8666460 | 14017739 | 4 | 30087466 | 40523834 |
| 1 | 359973 | 2826449 | 4 | 24704582 | 29206939 |
| 1 | 12094647 | 12676081 | 4 | 42831408 | 43911396 |
| 1 | 12293132 | 12500531 | 4 | 40588465 | 41503016 |
| 1 | 12233790 | 12280498 | 4 | 42165724 | 42373860 |
| 2 | 7999654 | 9110949 | 2 | 4224848 | 5625944 |
| 2 | 10929431 | 11237284 | 2 | 12038264 | 12900212 |
| 2 | 11260392 | 11512518 | 2 | 14000663 | 14325542 |
| 2 | 11958403 | 13338874 | 2 | 15733364 | 17813551 |
| 2 | 11552150 | 13910537 | 2 | 6017075 | 12033933 |
| 2 | 8834905 | 9131821 | 4 | 8110353 | 9886682 |
| 2 | 8787489 | 8837599 | 4 | 9544971 | 9732269 |
| 2 | 7268584 | 7966362 | 4 | 12953057 | 13570334 |
| 2 | 8976282 | 10913835 | 4 | 4292581 | 11192455 |
| 2 | 8512797 | 8767572 | 4 | 198622 | 1314837 |
| 2 | 8911312 | 9008149 | 4 | 2211188 | 2322502 |
| 2 | 13913258 | 14022952 | 4 | 11201127 | 11528901 |
| 2 | 10312728 | 10432051 | 4 | 1319560 | 1398290 |
| 2 | 14184276 | 14285240 | 4 | 12030284 | 12201931 |
| 2 | 14339213 | 14385948 | 4 | 12803310 | 12869632 |
| 2 | 14115374 | 14169337 | 4 | 12528395 | 12735463 |
| 2 | 14490 | 7008521 | 5 | 235042 | 21651067 |
| 2 | 7023641 | 7239531 | 5 | 14304402 | 17229375 |
| 3 | 79222 | 5052348 | 5 | 32591660 | 39048943 |
| 3 | 6299618 | 14303370 | 5 | 38930858 | 43639650 |
| 3 | 27144 | 1274802 | 5 | 26910056 | 29561462 |
| 4 | 11000514 | 13635366 | 2 | 29210091 | 33438038 |
| 4 | 7337778 | 11376941 | 2 | 37991307 | 40885670 |
| 4 | 5426131 | 5545754 | 2 | 37931095 | 38309427 |
| 4 | 3577950 | 3741224 | 2 | 37803680 | 37926686 |
| 4 | 319462 | 1744287 | 2 | 34642808 | 36986752 |
| 4 | 1914940 | 3263793 | 2 | 37072566 | 38852497 |
| 4 | 213107 | 317540 | 2 | 34459587 | 34625887 |
| 4 | 142996 | 190660 | 2 | 34192929 | 34306967 |
| 5 | 12929586 | 13254788 | 2 | 33459312 | 34138975 |
| 5 | 11775608 | 13003276 | 2 | 26940284 | 29189849 |
| 5 | 11306660 | 11768069 | 2 | 23753302 | 24718805 |
| 5 | 332362 | 1784009 | 3 | 49142220 | 51724044 |
| 5 | 90464 | 4134568 | 4 | 12803630 | 22397121 |
| 5 | 37612 | 230146 | 4 | 29807983 | 31297385 |
| 5 | 11528601 | 11683773 | 4 | 13581124 | 13886368 |
| 5 | 9162613 | 11291951 | 5 | 48426257 | 54179416 |
| 5 | 5750861 | 6975518 | 5 | 44411639 | 46434257 |
| 5 | 8906078 | 9151684 | 5 | 47015822 | 47261551 |
| 5 | 3959727 | 4508345 | 5 | 47406591 | 48404276 |
| 5 | 8289681 | 8888534 | 5 | 46350917 | 46813061 |
| 5 | 7055974 | 7986974 | 5 | 44143621 | 46286297 |
| 5 | 6736347 | 6830307 | 5 | 46955866 | 47011727 |
| 5 | 215266 | 2526380 | 6 | 62266276 | 65532391 |
| 6 | 12665565 | 12783255 | 2 | 43659754 | 43954101 |
| 6 | 56699 | 12660167 | 2 | 40524165 | 74700688 |
| 7 | 12204 | 11598106 | 6 | 2141059 | 32014494 |
| 8 | 10483 | 1835171 | 1 | 50269267 | 56114802 |
| 8 | 4849503 | 7242653 | 1 | 63401956 | 66398074 |
| 8 | 1830803 | 4641405 | 1 | 64684971 | 70832215 |
| 8 | 8150180 | 8383009 | 1 | 63267803 | 64381904 |
| 8 | 7413292 | 8054038 | 1 | 64064842 | 64259577 |
| 8 | 9126679 | 9777594 | 2 | 60984326 | 61613558 |
| 8 | 8405811 | 11256722 | 5 | 54186233 | 64025214 |
| 8 | 7723378 | 8125699 | 5 | 77447949 | 78052143 |
| 9 | 8832855 | 11257293 | 1 | 41687197 | 42944479 |
| 9 | 140956 | 269850 | 1 | 37168979 | 37445955 |
| 9 | 291145 | 9326323 | 1 | 27028971 | 41652557 |
| 10 | 5943186 | 10840555 | 6 | 41116401 | 52839742 |
| 10 | 122278 | 7810321 | 6 | 29413796 | 39548846 |
| 10 | 10756805 | 11052592 | 6 | 53709356 | 54367475 |
| 11 | 5394936 | 10692277 | 3 | 63207092 | 74523826 |
| 11 | 3410057 | 6094026 | 4 | 29928099 | 32748334 |
| 11 | 7209437 | 8101900 | 4 | 47566424 | 50981052 |
| 11 | 4062020 | 4504109 | 4 | 21445872 | 22401488 |
| 11 | 192880 | 2229452 | 4 | 44258164 | 46999064 |
| 11 | 8166596 | 11000846 | 4 | 22433661 | 29803767 |
| 11 | 7317021 | 7436411 | 4 | 56327500 | 56503390 |
| 11 | 7032696 | 7473377 | 4 | 57826318 | 59214129 |
| 11 | 6848299 | 7020873 | 4 | 34467566 | 34720837 |
| 11 | 6101077 | 6781651 | 4 | 32772382 | 34101216 |
| 11 | 6347286 | 6463921 | 4 | 34223055 | 34449008 |
| 12 | 55549 | 10651554 | 3 | 48188235 | 56853362 |
| 12 | 10251385 | 10587802 | 3 | 40361936 | 41139576 |
| 12 | 10654429 | 10826624 | 3 | 41140532 | 41820868 |
| 12 | 10225044 | 10553461 | 4 | 15593015 | 16866751 |
| 12 | 9729255 | 10780133 | 6 | 60830861 | 65601681 |
| 13 | 18100 | 3158839 | 1 | 56968706 | 66119576 |
| 13 | 1170242 | 1792460 | 1 | 74723398 | 75387382 |
| 13 | 3169326 | 3209736 | 1 | 61830310 | 61877603 |
| 13 | 2856704 | 6555343 | 1 | 71600637 | 75691791 |
| 13 | 372552 | 1226860 | 3 | 80922027 | 82644405 |
| 13 | 5805435 | 10477793 | 5 | 63090542 | 67917770 |
| 13 | 1023120 | 1345524 | 5 | 78168123 | 78515016 |
| 13 | 2545847 | 2753534 | 5 | 76821257 | 77435931 |
| 13 | 2796189 | 2852182 | 5 | 67921616 | 67973405 |
| 13 | 2043659 | 2092191 | 5 | 68763320 | 68826188 |
| 13 | 10339968 | 10438558 | 5 | 77819511 | 78144189 |
| 13 | 1774135 | 2016713 | 5 | 69371216 | 70184099 |
| 13 | 2371448 | 2536344 | 5 | 70215176 | 70482705 |
| 13 | 8204125 | 8928764 | 6 | 5780433 | 6500573 |
| 13 | 213669 | 816094 | 6 | 31851238 | 33142686 |
| 13 | 1590130 | 2374270 | 6 | 36712190 | 38152077 |
| 14 | 51508 | 4707248 | 1 | 2206837 | 11539468 |
| 14 | 7916567 | 8334659 | 1 | 36571036 | 37210484 |
| 14 | 6947042 | 7342120 | 1 | 24530289 | 26288997 |
| 14 | 5503388 | 6930468 | 1 | 17762939 | 19214086 |
| 14 | 8430276 | 8719711 | 1 | 11554265 | 13508709 |
| 14 | 7354360 | 10398935 | 1 | 20067174 | 27270539 |
| 14 | 9054306 | 9309721 | 1 | 16812313 | 17679531 |
| 14 | 8699711 | 9174423 | 5 | 25971431 | 26403506 |
| 15 | 566567 | 9896257 | 5 | 18675726 | 32575252 |
| 15 | 27749 | 4498029 | 5 | 23620505 | 29524166 |
| 15 | 8537137 | 10049083 | 5 | 25202396 | 32272251 |
| 15 | 6811992 | 7012002 | 5 | 15001436 | 15393055 |
| 16 | 441044 | 9736669 | 1 | 42611870 | 51404451 |
| 16 | 111293 | 183721 | 1 | 57245371 | 57346522 |
| 16 | 281605 | 1287513 | 1 | 56471918 | 58940483 |
| 16 | 24674 | 91588 | 1 | 56221516 | 56314036 |
| 16 | 212379 | 275809 | 1 | 56327650 | 56468170 |
| 17 | 54158 | 5265192 | 3 | 15660648 | 26438471 |
| 17 | 5611721 | 7618926 | 3 | 24472283 | 25658194 |
| 17 | 7440709 | 9240818 | 3 | 4383065 | 7736606 |
| 17 | 65687 | 629379 | 3 | 4331114 | 5967943 |
| 17 | 8287179 | 8648928 | 3 | 17682529 | 18084098 |
| 17 | 8944792 | 9598144 | 3 | 11892442 | 14529071 |
| 17 | 8651095 | 8815973 | 3 | 15581622 | 16248281 |
| 18 | 437536 | 2072472 | 3 | 80530283 | 82631536 |
| 18 | 39432 | 458984 | 3 | 75212573 | 76558228 |
| 18 | 5972559 | 9604378 | 6 | 67168275 | 76989589 |
| 18 | 2063417 | 4472133 | 6 | 82751148 | 84066213 |
| 18 | 3115209 | 4792432 | 6 | 71198605 | 72260985 |
| 18 | 7964222 | 8082089 | 6 | 82329533 | 82558149 |
| 18 | 8087692 | 8114236 | 6 | 82694279 | 82731526 |
| 19 | 75541 | 1051156 | 2 | 12915513 | 13906372 |
| 19 | 1061412 | 2541977 | 2 | 19631313 | 20676669 |
| 19 | 3162899 | 3481582 | 2 | 23632940 | 23721895 |
| 19 | 3586369 | 9249752 | 2 | 17869520 | 26934600 |
| 20 | 1257992 | 1774944 | 1 | 18370986 | 19797222 |
| 20 | 2055663 | 3343537 | 3 | 42907707 | 45830344 |
| 20 | 8379091 | 8477409 | 3 | 35410519 | 35567841 |
| 20 | 8471462 | 9083663 | 3 | 33829969 | 35397023 |
| 20 | 3421414 | 6346488 | 3 | 36206142 | 38884253 |
| 20 | 7044686 | 8368904 | 3 | 38974714 | 40326627 |
| 20 | 6388919 | 6802993 | 3 | 37013794 | 37536466 |
| 20 | 877364 | 996182 | 3 | 47306425 | 47548458 |
| 20 | 23000 | 2007100 | 3 | 41838156 | 45881483 |
| 20 | 9149727 | 9202636 | 3 | 11705871 | 11875903 |
| 20 | 6879827 | 6980210 | 3 | 48092288 | 48175010 |
| 20 | 583666 | 733059 | 3 | 35836505 | 36130630 |
| 21 | 851706 | 1571973 | 4 | 15488480 | 17273259 |
| 21 | 19744 | 6066179 | 6 | 52954241 | 67726371 |
| 21 | 5592418 | 6618700 | 6 | 54766857 | 55205196 |
| 21 | 6247286 | 7129442 | 6 | 57493146 | 57930594 |

